# Loss of TDP-43 induces synaptic dysfunction that is rescued by *UNC13A* splice-switching ASOs

**DOI:** 10.1101/2024.06.20.599684

**Authors:** Matthew J. Keuss, Peter Harley, Eugeni Ryadnov, Rachel E. Jackson, Matteo Zanovello, Oscar G. Wilkins, Simone Barattucci, Puja R. Mehta, Marcio G. Oliveira, Joanna E. Parkes, Aparna Sinha, Andrés F. Correa-Sánchez, Peter L. Oliver, Elizabeth M.C. Fisher, Giampietro Schiavo, Mala Shah, Juan Burrone, Pietro Fratta

## Abstract

TDP-43 loss of function induces multiple splicing changes, including a cryptic exon in the amyotrophic lateral sclerosis and fronto-temporal lobar degeneration risk gene *UNC13A*, leading to nonsense-mediated decay of *UNC13A* transcripts and loss of protein. UNC13A is an active zone protein with an integral role in coordinating pre-synaptic function. Here, we show TDP-43 depletion induces a severe reduction in synaptic transmission, leading to an asynchronous pattern of network activity. We demonstrate that these deficits are largely driven by a single cryptic exon in *UNC13A*. Antisense oligonucleotides targeting the *UNC13A* cryptic exon robustly rescue UNC13A protein levels and restore normal synaptic function, providing a potential new therapeutic approach for ALS and other TDP-43-related disorders.

## Main

Amyotrophic lateral sclerosis (ALS) is a rapidly progressive and fatal neurodegenerative condition. Recently, antisense oligonucleotides (ASOs) targeting *SOD1* and *FUS* have shown early promise in clinical trials^1,2^, but these approaches only benefit patients with *SOD1* and *FUS* mutations – less than 3% of ALS patients - leaving urgent need for therapies for the remaining ∼97% of ALS patients. A common feature of all non-FUS and -SOD1 ALS is the aggregation of TAR DNA binding protein-43 (TDP-43) in neurons^3,4^, a finding also observed in approximately 50% of both frontotemporal dementia (FTD) and Alzheimer’s disease (AD)^4,5^. However no therapies that directly target TDP-43 are available. TDP-43 is an RNA-binding protein that is normally located in the nucleus and has crucial roles in the regulation of RNA splicing^6,7^. Cytoplasmic mislocalization of TDP-43 induces a nuclear loss of function (LoF) resulting in cryptic exon inclusion, cryptic splice-polyadenylation or exon skipping in some mature mRNA transcripts^8^. These events occur in hundreds of transcripts, including genes with crucial neuronal functions, such as *STMN2*, *UNC13A*, and *KCNQ2* (ref. 9-12). The cryptic splicing-induced loss of *STMN2* was shown to affect axonal regeneration in vitro, leading to the development of splice-correcting ASOs as a therapeutic approach for ALS^13,14^.

Genetic evidence implicates loss of *UNC13A* in contributing to disease progression in ALS: multiple genome-wide association studies (GWAS) have shown that single nucleotide polymorphisms (SNPs) located within and proximal to the *UNC13A* cryptic exon increase the risk of developing ALS and accelerate disease progression^15–17^. Under normal physiological conditions, these risk SNPs do not induce inclusion of the *UNC13A* cryptic exon, but upon TDP-43 LoF, the presence of risk SNPs exacerbates the cryptic exon inclusion leading to loss of functional RNA transcripts and proteins. Whilst this association provides strong genetic relevance for this event in ALS pathogenesis, the impact of TDP-43 LoF and the *UNC13A* cryptic exon on synaptic function are still unknown. UNC13A is a presynaptic protein that is required at the active zone to properly dock and prime synaptic vesicles for fusion with the plasma membrane. Complete genetic deletion of *Unc13a* in mice leads to a strong reduction in spontaneous and evoked glutamatergic excitatory synaptic transmission^18–20^. These results highlight the importance of UNC13A in neurotransmitter release and open the possibility that synapse function may be compromised in ALS. If so, correction of *UNC13A* splicing could be a viable approach to recover normal neuronal function.

Here, we report that TDP-43 loss in mature human iPSC-derived neurons induces loss of pre-synaptic UNC13A and a drastic reduction in AMPA-mediated synaptic transmission, leading to a disordered and asynchronous pattern of network activity. Remarkably, CRISPR/Cas9 deletion of the *UNC13A* cryptic exon rescues these deficits demonstrating this is a crucial alteration driving synapse dysfunction in ALS. Furthermore, ASOs targeting the cryptic exon in *UNC13A* rescue mRNA splicing, protein levels, UNC13A pre-synaptic localisation and restore synaptic transmission. Thus, ASOs targeting the *UNC13A* cryptic exon represent a novel therapeutic strategy in ALS-FTD.

### Loss of TDP-43 in neurons leads to depletion of synaptic UNC13A

TDP-43 pathology and ALS typically start after nervous system development and decades of normal function. TDP-43 plays important roles in development and therefore to avoid interfering with neuronal maturation by reducing TDP-43 from the iPSC stage, we used CRISPR/Cas12 gene editing to generate a cell line where both endogenous copies of TDP-43 harbour a HaloTag to allow for proteolysis targeting chimera (PROTAC)-mediated knockdown of TDP-43 in mature neurons by the Cul2/VHL ubiquitin E3-ligase (Fig. 1a and Extended Data Fig. 1a)^21^. TDP-43 is degraded by HaloPROTAC in a dose-dependent manner and in a process blocked by proteasome inhibition or the NEDD8 E1 inhibitor MLN4924, which prevents the activation of Cullin-RING ubiquitin E3-ligases (Extended Data Fig.1b-d). HaloTag TDP-43 iPSCs were differentiated into mature neurons (Halo-iNeurons) where TDP-43 correctly localised to the nucleus (Fig. 1b). Tagging of TDP-43 can alter its function and, whilst the HaloTag partially affected TDP-43 function (Extended Data Fig. 1e), *UNC13A* cryptic splicing was not detected in basal conditions, and only upon HaloPROTAC knockdown of TDP-43 were *UNC13A* cryptic exons and mRNA degradation observed (Fig. 1c,d and Extended Data Fig. 1f), similarly to CRISPRi-mediated knockdown (Extended Data Fig. 1g-k). Loss of *UNC13A* mRNA led to a time-dependent decrease in UNC13A protein whereby less than 5% of protein was left after 10 days of HaloPROTAC treatment (Fig. 1e,f). Immunofluorescence staining revealed a loss of synaptic UNC13A staining at synapsin-positive puncta (Fig. 1g,h). HaloPROTAC-mediated knockdown of TDP-43 in mature neurons did not lead to alterations in the RNA levels of NMDA and AMPA glutamate receptor subunits as assessed by RT-qPCR (Extended Data Fig. 1l).

**Fig. 1.**
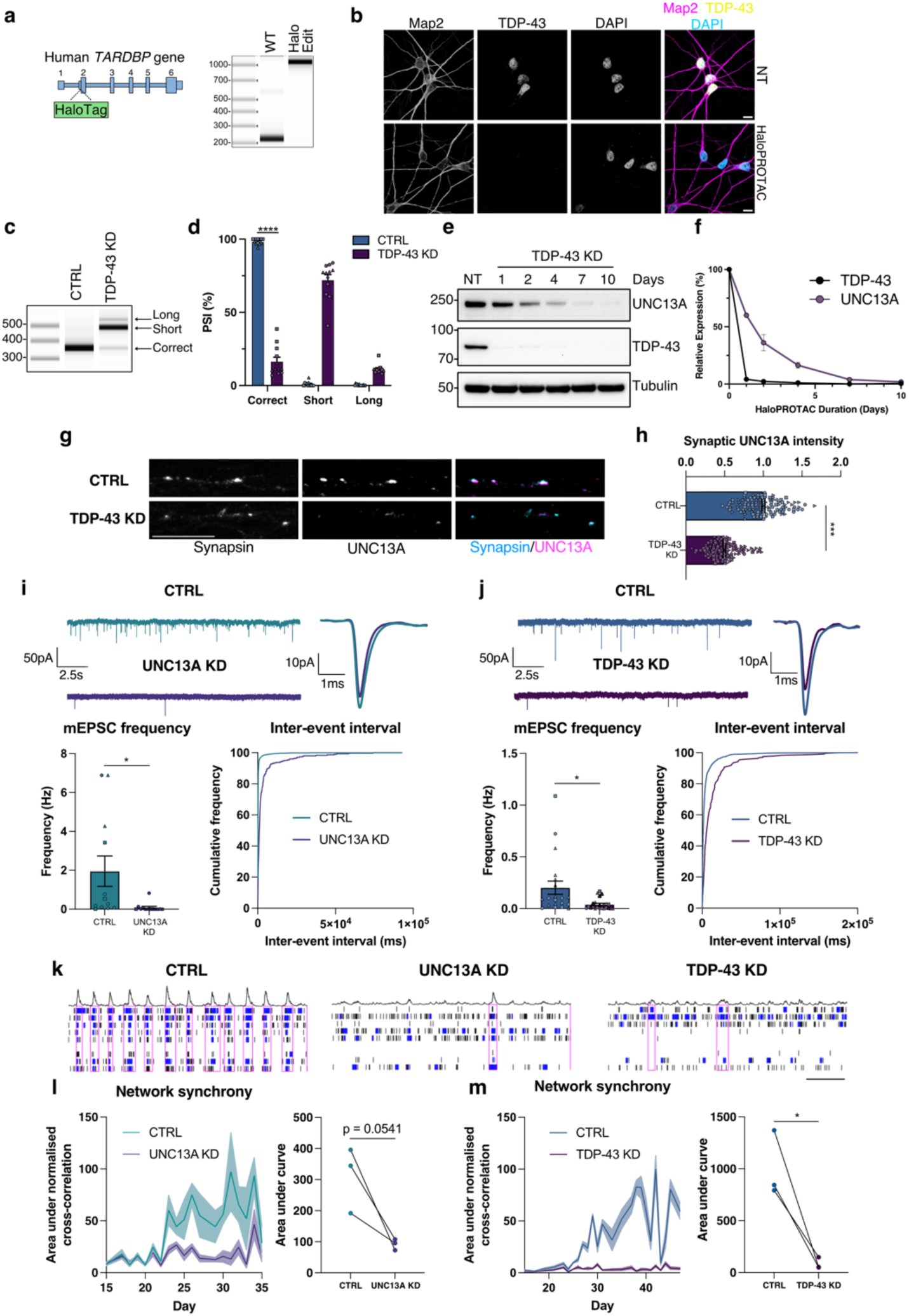
UNC13A and TDP-43 knockdown impair synaptic function. **a**, Schematic representation of HaloTag edit at the beginning of exon 2 of *TARDBP* and PCR from genomic DNA indicating successful edit **b**, Immunofluorescence analysis of TDP-43 localization in HaloTDP iNeurons revealed nuclear localised TDP-43. Scale bar = 10 µm. **c**, RT-PCR analysis of *UNC13A* splicing after TDP-43 knockdown. **d**, Quantification of results in (**c**) *n*=12 biological replicates from 3 experiments. **e**, Western blot analysis of HaloTDP iNeurons treated with 30 nM HaloPROTAC-E for the indicated amount of time. **f**, Quantification of western blot in (**e**) *n*=2 biological replicates. **g**, Immunofluorescence labelling of synapsin and UNC13A pre-synaptic terminals in 4-week-old HaloTDP iNeurons grown on rat astrocytes following 2-weeks of TDP-43 depletion. Scale bar = 10 µm. **h**, Quantification of UNC13A intensity at synapsin positive synaptic terminals. Control n=116, TDP-43 knockdown *n*=115 synapses from 3 experiments. **i**, Miniature excitatory postsynaptic currents (mEPSCs) from control (*n*=12) and UNC13A knockdown (*n*=13) iNeurons pooled from 3 experiments. Representative traces and average traces shown. Quantification of mEPSC frequency and inter-event interval cumulative distributions. **j**, mEPSCs from control (*n*=20) and TDP-43 knockdown (*n*=22) HaloTDP iNeurons pooled from 3 experiments. Representative traces and average traces shown. Quantification of mEPSC frequency and inter-event interval cumulative distributions. **k**, Representative raster plots from D36 multi-electrode array recordings from control, UNC13A and TDP-43 knockdown conditions (30s window shown). Electrode bursts in blue and network bursts in pink. **l**, Time-course of network synchrony measured by the area under the normalised electrode cross correlation in control and UNC13A knockdown iNeurons. UNC13A normalised to control *n*=18 wells from 3 experiments. **m**, Time-course of network synchrony in control and TDP-43 knockdown iNeurons. TDP-43 normalised to control, *n*=18 wells from 3 experiments. Graphs for (**d**) (**h**) (**i**) and (**j**) represent mean ± s.e.m and independent experiments are denoted by circle, square or triangle markers. Statistics for (**d**) (**h**) (**i**) and (**j**) are two-sided Student’s *t* test. Statistics for L and M are ratio paired *t* tests. \**P* < 0.05; ** *P* <0.01; *** *P* <0.001; **** *P* < 0.0001; ns (not significant).

### Loss of TDP-43 in neurons impairs synaptic function

Previous investigations of the physiological functions of UNC13A have been conducted using primary rodent neurons. We therefore sought to assess the impact of *UNC13A* loss in human iPSC-derived neurons. Whole-cell patch clamp recordings showed that depletion of *UNC13A* via CRISPRi impaired synaptic transmission as measured by a reduction in the frequency of miniature excitatory postsynaptic currents (mEPSC) (Fig. 1i) in line with previously published work^19,20^. Intriguingly, HaloPROTAC-mediated TDP-43 depletion also impaired synaptic transmission in a comparable manner (Fig. 1j). CNQX treatment blocked mEPSCs, confirming that synaptic events were mediated primarily via fast AMPA receptor-mediated glutamatergic transmissions (Extended Data Fig. 2a). While UNC13A depletion had no effect on mEPSC amplitude, usually indicating postsynaptic alterations (Extended Data Fig. 2b), TDP-43 depletion led to a modest reduction in amplitude (Extended Data Fig. 2c), suggesting additional postsynaptic deficits independent of direct UNC13A depletion. Passive membrane properties remained comparable between control and UNC13A/TDP-43 depleted neurons (Extended Data Fig. 2d,e).

To investigate the effect of UNC13A and TDP-43 loss on network activity, we carried out micro-electrode array (MEA) recordings. The number of active electrodes remained consistent (Extended Data Fig. 2g,h) and phase contrast images showed comparable cell coverage between conditions (Extended Data Fig. 2i) ruling out issues with electrode coverage and cell viability as confounding factors. We found that UNC13A depletion induced a disordered and asynchronous pattern of network activity, measured by a reduction in the normalised electrode cross-correlations (Fig. 1k,l). TDP-43 depletion also induced a very similar deficit in network synchrony (Fig. 1k,m). CNQX treatment confirmed that this readout of network synchrony was highly dependent on fast AMPA receptor-mediated synaptic transmission, (Extended Data Fig. 2f). Other metrics such as mean firing rate were only minimally affected by UNC13A and TDP-43 depletion (Extended Data Fig. 2g,h). Thus, we show that UNC13A loss substantially impairs fast glutamatergic synaptic transmission and synchronised network activity in human neurons, mirroring a comparable loss of synaptic function and network activity induced by TDP-43 depletion.

### Genomic deletion of *UNC13A* cryptic exon prevents UNC13A protein loss

Knockdown of TDP-43 leads to cryptic exon inclusion within *UNC13A* intron 20 along with an independent downstream intron retention event at intron 31 and additional lower frequency splicing abnormalities throughout the *UNC13A* mRNA transcript^11,12^. To determine the degree to which protein loss is dependent on the *UNC13A* intron 20 cryptic exon, we performed CRISPR/Cas9 gene editing to remove the cryptic exon from the genome of SH-SY5Y cells (Fig. 2a and Extended Data Fig. 3a,b). Using two sgRNAs we made a homozygous deletion of 233 nucleotides which contain both splice acceptors and the common donor splice site of the cryptic exon. In WT cells containing the full *UNC13A* gene there was robust induction of *UNC13A* mis-splicing after TDP-43 depletion as assessed by RT-PCR (Fig. 2b,c). While in cells with genomic deletion of the *UNC13A* cryptic exon, no cryptic splicing was detected at the *UNC13A* exon 20-21 junction (Fig. 2b,c). Cryptic exon deletion rescued correctly spliced levels of *UNC13A* mRNA as determined by RT-qPCR and led to ∼75% rescue of levels of UNC13A protein following TDP-43 knockdown (Fig. 2c-f). The residual TDP-43-dependent loss of UNC13A protein after cryptic exon deletion was likely due to other *UNC13A* splicing alterations as the intron retention event at exon 31-32 that is increased after TDP-43 knockdown^11^. We subsequently deleted the *UNC13A* cryptic exon in iPSCs (Extended Data Fig. 3c,d) and detected a similar rescue in *UNC13A* RNA and protein following CRISPRi-mediated knockdown of TDP-43 (Extended Data Fig. 3e-j). In the *UNC13A* cryptic exon deletion cell line, we performed additional CRISPR/Cas12 gene editing to add a HaloTag in the *TARDBP* gene (Extended Data Fig. 3k,l). Similar to results in SH-SY5Y cells, genomic deletion of the *UNC13A* cryptic exon in Halo-iNeurons rescues *UNC13A* splicing at the exon 20-21 junction and restores mRNA and protein levels (Fig. 2g-k) demonstrating this splicing event is the main driver of TDP-43-dependent UNC13A loss.

**Fig. 2.**
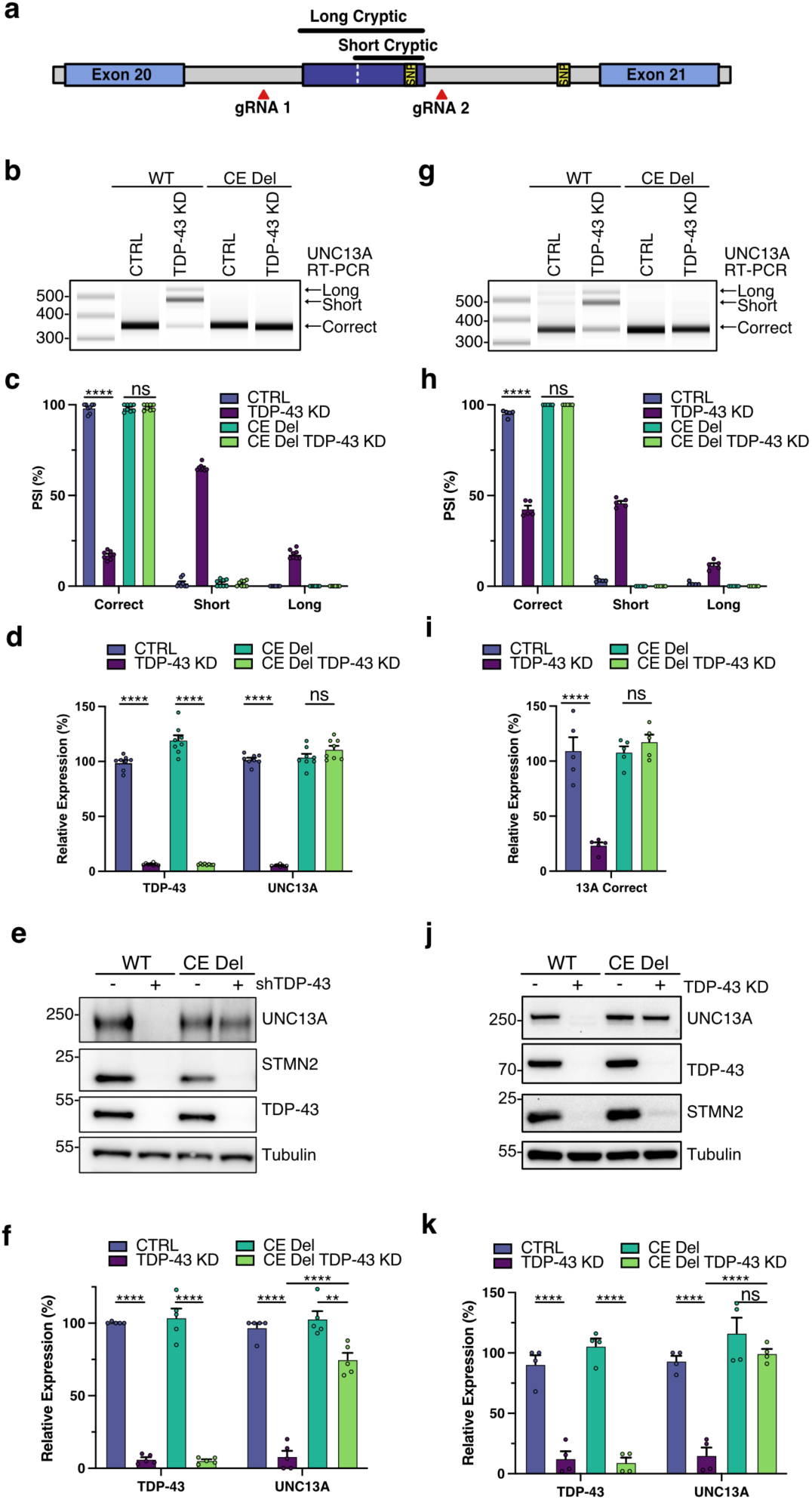
Genomic deletion of *UNC13A* CE rescues protein levels following TDP-43 knockdown. **a**, Schematic representation of genomic edit of *UNC13A* cryptic exon. **b-f**, Analysis of control and TDP-43 KD in WT and CE Del SH-SY5Y cells. (**b**) RT-PCR analysis of *UNC13A* splicing indicates CE deletion rescues splicing at the *UNC13A* exon 20-21 junction after TDP-43 KD. (**c**) Quantification of results in (B) *n*=8 biological replicates from 3 experiments. (**d**) RT-qPCR analysis of TDP-43, correctly spliced *UNC13A* shows CE deletion rescues *UNC13A* RNA levels after TDP-43 KD. *n* = 8 biological replicates from 3 experiments. (**e**) Western blot analysis of protein lysates shows CE deletion rescues UNC13A protein levels after TDP-43 KD. (**f**) Quantification of results in (**e**) *n*=5 biological replicates from 3 experiments. (**g-k**) Analysis of control and TDP-43 knockdown in WT and CE Del Halo-TDP iNeurons on DIV 28 following 14 days of HaloProtac treatment. (**g**) RT-PCR analysis of *UNC13A* splicing indicates CE deletion rescues splicing at *UNC13A* exon 20-21 junction after TDP-43 KD. (**h**) Quantification of results in (**g**) *n*=5 biological replicates from 2 experiments. (**i**) RT-qPCR analysis of correctly spliced *UNC13A* shows CE deletion rescues *UNC13A* RNA levels after TDP-43 KD. *n*=5 biological replicates from 2 experiments. (**j**) Western blot analysis of protein lysates indicates CE deletion rescues UNC13A protein levels after TDP-43 KD. (**k**) Quantification of results in (**j**) *n*=4 biological replicates from 2 experiments. Graphs for (**c**) (**d**) (**f**) (**h**) (**i**) and (**k**) represent mean ± s.e.m. One-way ANOVA with Tukey multiple comparison test. \**P* < 0.05; ** *P* <0.01; *** *P* <0.001; **** *P* < 0.0001; ns (not significant).

### Genomic deletion of *UNC13A* cryptic exon rescues synaptic function

TDP-43 knockdown yields similar synaptic defects as those induced by direct depletion of *UNC13A*. However, TDP-43 depletion also induces widespread splicing defects in other synaptic genes, and therefore the degree to which *UNC13A* loss is responsible for the TDP-43 knockdown phenotype is unknown. To address this, we sought to determine to what extent the *UNC13A* intron 20 cryptic exon (CE) contributes to impaired synaptic function following loss of TDP-43. CRISPR/Cas9 genomic deletion of the *UNC13A* cryptic exon rescued pre-synaptic UNC13A back to normal levels following depletion of TDP-43 (Fig. 3a,b and Extended Data Fig. 4a). In conjunction with a rescue of synaptic UNC13A, deletion of the *UNC13A* cryptic exon rescued synaptic transmission following TDP-43 depletion. mEPSC frequency was rescued back to near normal levels, while mEPSC amplitude was not, suggesting a specific rescue in pre-synaptic function mediated by the rescue of UNC13A (Fig. 3c,d and Extended Data Fig 4b). Passive membrane properties remained unchanged between conditions (Extended Data Fig. 4c).

**Fig. 3.**
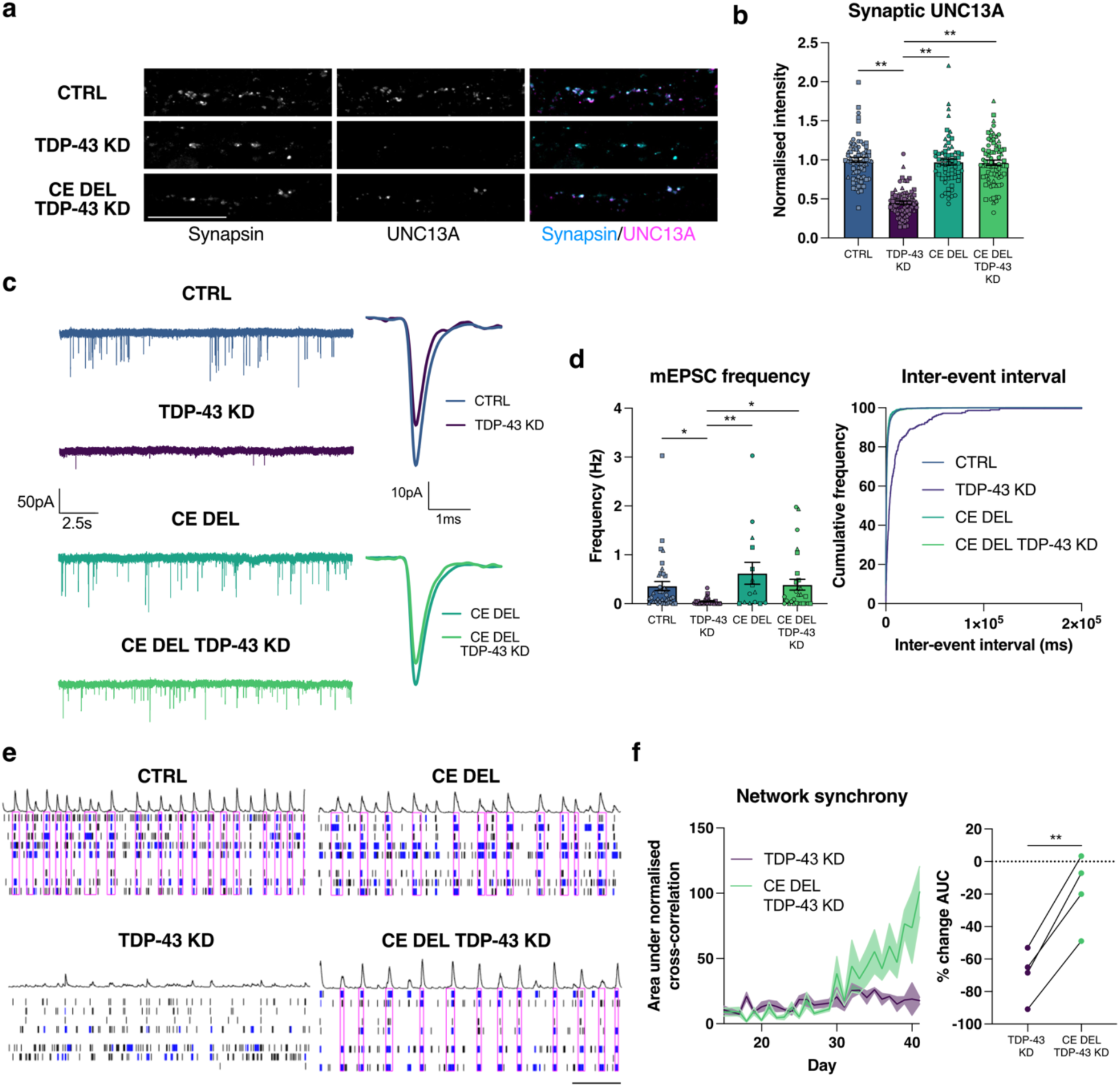
Genomic deletion of *UNC13A* CE rescues synaptic function following TDP-43 knockdown. **a,** Immunofluorescence labelling of synapsin and UNC13A pre-synaptic terminals in 4-week old HaloTDP and *UNC13A* CE Del iNeurons grown on rat astrocytes following 2-weeks of TDP-43 knockdown. Scale bar = 10 µm. **b**, Quantification of UNC13A intensity at synapsin positive synaptic terminals. Control *n*=71, TDP-43 knockdown *n*=71, CE Del *n*=66, CE Del TDP-43 knockdown *n*=75 synapses from 3 experiments. **c,** Representative mEPSC traces and average amplitude traces. **d,** Quantification of mEPSC frequency and inter-event interval cumulative distributions from control *n*=17, TDP-43 knockdown *n*=22, CE Del n=15, CE Del TDP-43 knockdown *n*=28 iNeurons pooled from 3 experiments. **e**, Representative raster plots from D39 multi-electrode array recordings from control, TDP-43 knockdown, CE Del, and CE Del TDP-43 knockdown conditions (30s window shown). Electrode bursts in blue and network bursts in pink. **f**, Time-course of network synchrony measured by the area under the normalised electrode cross correlation with TDP-43 knockdown conditions normalised to each control *n*=24 wells from 4 experiments. Graphs for (**b**) (**d**) (**f**) represent mean ± s.e.m. Statistics for (**b**) and (**d**) are one-way ANOVAs with Dunnet’s multiple comparisons test, statistics for (**f**) is a paired *t* test. \**P* < 0.05; ** *P* <0.01.

At a network level there was a significant and long-term rescue in the disordered and asynchronous pattern of network activity induced by TDP-43 depletion. Deletion of the *UNC13A* cryptic exon restored network synchrony, measured by cross-correlations between electrodes, to near normal levels, (Fig. 3e,f). Taken together, the data show that TDP-43 depletion results in impaired synaptic transmission, largely via the *UNC13A* cryptic exon. This loss in synaptic transmission manifests as a highly disordered and asynchronous pattern of network activity that can be rescued by directly deleting the *UNC13A* cryptic exon.

### Antisense oligonucleotides rescue *UNC13A* splicing

The rescue of UNC13A levels and restoration of synaptic signalling conferred by deletion of the *UNC13A* cryptic exon in the context of TDP-43 knockdown highlights the therapeutic potential of targeting this event in patients. We therefore sought to identify ASOs that can induce splice switching similar to those successful in targeting splicing events in *SMN2* and *STMN2* (ref. 13,22). Initially, we designed locked nucleic acids (LNA)-modified ASOs that targeted regions covering the cryptic exon acceptor and donor splice sites (Extended Data Fig. 5a). We screened the ASOs by transfection in SK-N-BE(2) cells and assessed their ability to restore *UNC13A* splicing at exon 20-21 by RT-PCR (Extended Data Fig. 5b) and identified multiple 21 nt ASOs that could restore *UNC13A* splicing. We confirmed splicing rescue and ability to restore *UNC13A* RNA and protein levels in SH-SY5Y cells (Extended Data Fig. 5c-g). To improve therapeutic tolerance, we redesigned the best performing ASOs to have full 2’-*O*-methoxyethyl ribose (MOE)-modified bases and tested their ability to rescue *UNC13A* splicing and restore protein levels in Halo-iNeurons (Fig. 4a-c and Extended Data Fig. 5h). We found these could restore splicing to >90% and protein levels to 30-50% of normal levels. Treatment with the M21-4 ASO led to increased pre-synaptic UNC13A protein expression at synapsin-positive puncta following TDP-43 depletion as compared to treatment with the control scrambled ASO (Fig. 4d). To identify ASOs that could increase protein rescue above 50% of normal levels we carried out a secondary screen by performing an ASO walk along the cryptic donor site with 18 nt MOE-based ASOs (Extended Data Fig. 6a). We screened the ability of the 18 nt ASOs to rescue splicing in SH-SY5Ys by RT-PCR (Extended Data Fig. 6b) and we confirmed that the best performing 18 nt ASOs could rescue *UNC13A* splicing and mRNA to >90%, and protein to >50% in both SH-SY5Y cells and Halo-iNeurons (Fig. 4e,f and Extended Data Fig. 6c-h). Overall, these results show that ASOs can rescue *UNC13A* cryptic exons and re-establish levels of UNC13A protein to above 50% of normal.

**Fig. 4.**
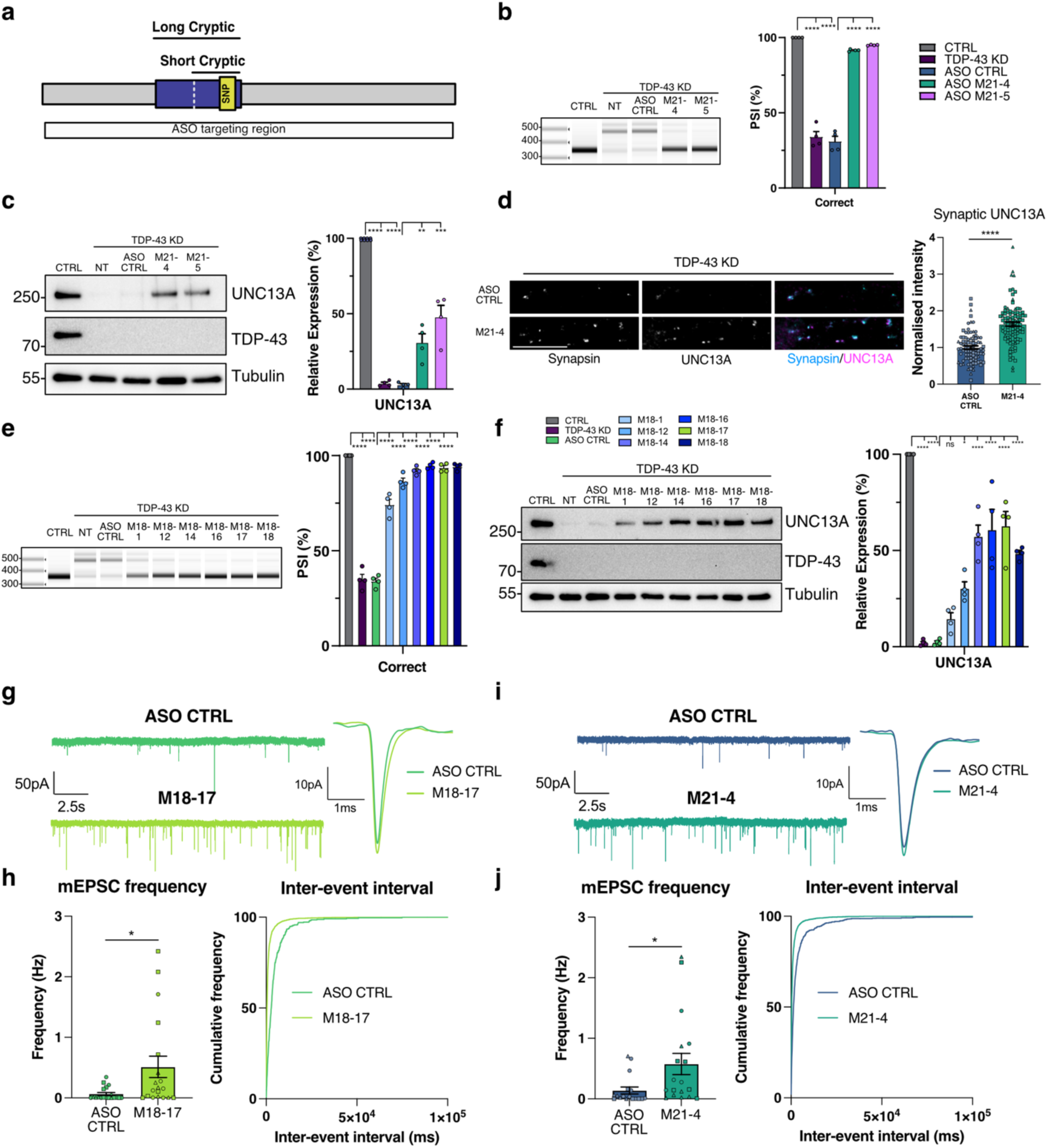
ASO mediated correction of *UNC13A* CE rescues synaptic function following TDP-43 knockdown. **a**, Schematic of *UNC13A* cryptic exon. **b**, RT-PCR analysis of *UNC13A* splicing in HaloTDP iNeurons shows 21nt ASOs prevent cryptic splicing after TDP-43 knockdown with quantification of results *n*=4 biological replicates from 2 experiments. **c**, Western blot analysis shows rescue of UNC13A protein following 21nt ASO treatment after TDP-43 KD with quantification. *n*=4 biological replicates from 2 experiments. **d**, Quantification of UNC13A intensity at synapsin positive synaptic terminals in TDP-43 knockdown iNeurons treated with MOE CTRL ASO (*n*=88) and M21-4 ASO (n=96). Scale bar = 10 µm. **e**, RT-PCR analysis of *UNC13A* splicing in HaloTDP iNeurons shows 18 nt ASOs prevent cryptic splicing after TDP-43 knockdown with quantification of results *n*=4 biological replicates from 2 experiments. **f**, Western blot analysis shows rescue of UNC13A protein following 18 nt ASO treatment after TDP-43 KD with quantification. *n*=4 biological replicates from 2 experiments. **g**, Representative mEPSCs traces and average mEPSCs traces from TDP-43 KD iNeurons treated with MOE CTRL ASO (n=17) and M21-4 ASO (*n*=18). **h**, Quantification of mEPSC frequency and inter-event interval cumulative distributions. **i,** Representative mEPSCs traces and average mEPSCs traces from TDP-43 KD iNeurons treated with CTRL ASO (*n*=19) and M18-17 ASO (*n*=19). **j**, Quantification of mEPSC frequency and inter-event interval cumulative distributions. Graphs for (**b**) (**c**) (**d**) (**e**) (**f**) (**h**) and (**j**) represent mean ± s.e.m. Statistics for (**b**) (**c**) (**e**) and (**f**) are One-way ANOVA with Tukey multiple comparison test. Statistics for (**d**) (**h**) and (**j**) are two-sided Student’s *t* test. \**P* < 0.05; ** *P* <0.01; *** *P* <0.001; **** *P* < 0.0001; ns (not significant).

### Antisense oligonucleotides rescue synaptic function

*Unc13a* heterozygous KO mice and individuals with heterozygous loss of function *UNC13A* mutations are viable and do not present with abnormal neurophysiology. Full recovery of *UNC13A* expression after TDP-43 LoF is therefore not required and recent work in primary mouse cultures has shown that clear deficits in synapse function are only observed when expression of *Unc13A* is below 30% (ref. 20). Treatment with M18-17 ASO, which rescues UNC134A levels to >50% of control, was able to rescue synaptic transmission by increasing mEPSC frequency to levels similar to those previously observed without TDP-43 loss (Fig. 4i,j). We then tested our M21-4 ASO, which rescues UNC13A protein levels to ∼30% of normal. Despite the lower level of protein rescue, treatment with M21-4 ASO rescued synaptic transmission following TDP-43 depletion as measured by an increase in mEPSC frequency (Fig. 4g,h). mEPSC amplitude was unaffected by treatment with ASOs (Extended Data Fig. 7a,b), suggesting that ASO treatment specifically rescues pre-synaptic deficits caused by the cryptic exon in *UNC13A*. Taken together these data show how therapeutic targeting of the *UNC13A* cryptic exon using an antisense oligonucleotide can rescue dysfunctional synaptic transmission caused by TDP-43 depletion.

## Discussion

TDP-43 cytoplasmic aggregation and nuclear loss of function (LoF) occur in approximately 97% of ALS patients^4^. Loss of nuclear TDP-43 leads to numerous changes in RNA processing, including the appearance of hundreds of novel exon inclusion, exon skipping, and polyadenylation events that are not observed under normal conditions, hence termed “cryptic” ^11,23^. Many such events result in the loss of transcripts and encoded proteins. However, one cryptic exon in intron 20 of the *UNC13A* gene has attracted significant interest, as SNPs accounting for the second strongest ALS GWAS signal are located within this intron^15–17^.

TDP-43 loss induces the *UNC13A* cryptic exon in alleles with either the risk or reference SNPs, but the risk SNPs enhance cryptic splicing, leading to decreased transcript and protein levels. The risk SNPs are associated with shorter survival. Therefore, regardless of how *UNC13A* loss contributes to disease pathogenesis, genetic data strongly support targeting this event to modify disease progression. Importantly, our experiments were conducted in iPSC neurons heterozygous for the SNPs and demonstrate that ASOs can effectively target both alleles, thus potentially benefiting virtually all individuals with TDP-43 pathology, irrespective of their genotype.

TDP-43 has multiple binding sites within the *UNC13A* pre-mRNA, therefore its depletion can be associated with multiple splicing changes, such as the increase in intron 31 retention. Deleting the intron 20 event significantly rescues protein levels to >75% of control, further demonstrating that the intron 20 cryptic exon is the dominant driver of TDP-43-induced UNC13A loss, with other changes possibly accounting for the remaining ∼25%.

The mechanism and functional implications of the *UNC13A* genetic association, which is also replicated in an FTLD-TDP GWAS^24^, remain unknown. UNC13A protein is crucial for synaptic vesicle priming; *Unc13A* null mice die at birth from respiratory failure, and primary cortical cultures from these mice develop normally but have significantly reduced excitatory synaptic transmission. Data on human neurons is more limited^17^, and while ALS-linked mutations have been shown to cause excitability and synaptic deficits in human neurons^25–27^, it remains to be seen whether direct depletion of TDP-43 and associated alterations in RNA splicing also drive impaired synaptic function. Here, we confirm that in human iPSC-derived neurons, loss of UNC13A recapitulates the reduction in spontaneous synaptic release well-described in primary neurons, leading to a disordered and asynchronous pattern of network activity. Intriguingly, TDP-43 loss also mirrored these deficits in synaptic function and network activity.

TDP-43 loss affects many transcripts and proteins that can impact neuronal maturation. Therefore, we devised an experimental set-up in which TDP-43 loss is induced after neuronal maturation, thus circumventing maturation changes and better matching the series of events occurring in patients, where TDP-43 pathology and loss occur after decades of normal neuronal function. We observed that TDP-43 loss gives rise to both a synaptic spontaneous release deficit and loss of network synchrony comparable to those observed after direct *UNC13A* knockdown. TDP-43 targets numerous transcripts involved in neuronal transmission, by binding to long introns, UTRs, or by inducing cryptic exons. To assess the contribution of *UNC13A* to this synaptic phenotype, we selectively deleted the *UNC13A* intron 20 cryptic exon and showed that this could rescue both the spontaneous release and network synchrony deficits, concluding that loss of UNC13A is the main driver of this phenotype upon TDP-43 loss. Intriguingly, also *UNC13B* undergoes alternative splicing upon TDP-43 depletion, leading to protein loss^11^. Although work in primary cultures has highlighted the more prominent role for UNC13A in synaptic vesicle release^19^, consistent with our findings, further research will be needed to assess whether UNC13B also contributes to TDP-43 synaptic dysfunction. It is likely that other synaptic genes could play further roles and contribute to other phenotypes - intriguingly we observe amplitude changes upon TDP-43 loss, which are not present upon direct *UNC13A* loss, nor are rescued by deleting the *UNC13A* cryptic exon. Further studies will be needed to elucidate the genes or functional mechanisms underpinning such changes.

Although data assessing UNC13A protein levels in patient neurons are not available, transcriptomic data on post-mortem neurons with TDP-43 pathology show that the *UNC13A* cryptic exon can be included in over >90% of transcripts, which is thus predicted to greatly inhibit production of functional UNC13A protein^28^. These findings support the validity of using an iPSC-derived neuronal model where TDP-43 loss induces a near-total loss of UNC13A. Our results show that restoring 30% of UNC13A levels can rescue spontaneous vesicle release, and heterozygous carriers of LoF *UNC13A* mutations do not exhibit overt phenotypes. This suggests that splice-switching therapeutics that lead to such a rescue in patients could contribute to the recovery of synaptic function.

The limited knowledge about the role of *UNC13A* loss of function in human disease comes from a report of a patient with inherited mutation in the gene. In this case, an individual developed microcephaly and myasthenia upon biallelic truncating mutation in *UNC13A* and had cortical hyperexcitability upon EEG analysis; the patient died at 50 months of age of respiratory failure^17^. Similar to this patient, transcranial magnetic stimulation studies have detected cortical hyperexcitability at an early stage in ALS patients that worsens with disease progression^22^. Moreover, one of the few drugs that extend survival in ALS, riluzole, causes a reduction in cortical hyperexcitability, suggesting this phenotype may be relevant for the survival of affected people^23^. Cortical hyperexcitability has also been shown for individuals with mutations in *STX1B* and *STXBP1* (ref. 18,19), and *SNAP25B*^20^, other presynaptic proteins that regulate synaptic vesicle release. To try and dissect the cause of hyperexcitability in vivo, researchers developed a mouse model of *Stxbp1* haploinsufficiency. This showed that impaired synaptic output from upstream excitatory neurons decreases the probability of action potential firing in inhibitory neurons. This results in an impairment in microcircuit feedforward inhibition and an overall skew of the excitatory/inhibitory ratio, eventually causing the hyperexcitable state^21^.

The cellular events that lead to TDP-43 aggregation in ALS are unknown and it is therefore challenging to correct this directly, but aggregation is associated with a loss of TDP-43 nuclear function that produces numerous cryptic splicing events, which may represent more tractable therapeutic targets than targeting mislocalization directly. Splice modifying ASOs have been successfully developed to treat spinal muscular atrophy by correcting *SMN2* splicing^22,29^. This same strategy can be used to correct cryptic splicing in ALS, yet there is still a need to identify splicing events that have the strongest impact on neuronal function and hold the most promise of improving disease symptoms. Our findings, along with results from GWAS studies showing that modulation of the *UNC13A* cryptic exon impacts disease progression^16^, strongly support *UNC13A* as a key candidate for splice-modifying therapy development in ALS. TDP-43 LoF drives sufficient UNC13A reduction to impair synaptic transmission and network synchrony. Prevention of *UNC13A* mis-splicing via CRISPR/Cas9 gene editing or application of ASOs can rescue the majority of UNC13A protein and restore defects in the frequency of mEPSCs. Our ability to rescue UNC13A protein and function in TDP-43 knockdown neurons is an exciting therapeutic avenue for ALS/FTD.

## Materials and Methods

### Cell Culture

SH-SY5Y cells (ATCC) and SK-N-BE(2) (ATCC) with stable doxycycline inducible shRNA to *TARDBP* were previously described^11^ and were maintained in DMEM/F12 with Glutamax (Thermo) with 10% FBS (Thermo) and split with TrypLE Express (Thermo) when 80% confluent. The WTC11 iPSC line derived from a healthy human subject (GM25256) with stable integration of doxycycline-inducible expression of *NGN2* at the *AAVS1* locus and stable integration of dCas9-BFP-KRAB at the *CLYBL* locus was a gift from Michael Ward^30^ were maintained in E8 Flex Medium (Thermo) in Geltrex (Thermo) coated plates and passed with Versene solution (Thermo) or Accutase (Thermo) when 80% confluent. For induction of iPSCs to iNeurons, iPSCs were passaged with Accutase (Thermo) and plated into Geltrex-coated plates with induction media: DMEM/F12 with Glutamax (Thermo), 1x Non-essential amino acids (Thermo), 2 μg/ml doxycycline hyclate (Sigma), 2 μM XAV939 (Cayman Chemical), 10 μM SB431542 (Cayman Chemical) and 100 nM LDN-193189 (Cayman Chemical)^31,32^. Media was changed daily for three days. On the third day cells were replated to poly-L-ornithine coated plates for RNA and protein experiments or on poly-D-lysine and laminin coated 18 mm coverslips (Deckglaser) with primary rat astrocytes for electrophysiology and immunofluorescent experiments. Differentiated neurons were maintained in neuron media (BrainPhys (StemCell Technologies) with 1x N2 supplement (Thermo), 1x B27 supplement (Thermo), 20 ng/mL BDNF (Peprotech), 20 ng/ml GDNF (Peprotech), 1 mM dibutyrl cAMP (Sigma), 200 nM L-ascorbic acid (Sigma), and 1 μg/mL laminin (Thermo))^33^ and half media changes were performed twice per week.

### Antisense oligonucleotide synthesis

LNA based ASOs were purchased from IDT. 21 nt MOE ASOs were purchased from Dharmacon. A panel of 18 nucleotide antisense oligonucleotides (ASO) were designed to tile across the region of interest before being subjected to further selection based on potential sequence-driven off-target effects and secondary structure. The resulting panel of twenty four 2’MOE ASOs was generated by solid-phase synthesis using MerMade12 automated synthesizer using pre-packed CPG columns (1 µmol scale, 1000Å). Phosphoramidites (TheraPure by Thermo) were prepared at 0.08 M using acetonitrile. Cleavage and deprotection of oligonucleotides were performed using concentrated ammonia 6 hours at 55°C. After deprotection, samples were dried and resuspended in water. Purification of the oligonucleotides was performed by reversed-phase HPLC using 0.1 M triethylammonium acetate pH 7 and acetonitrile in an XBridge BEH C18 Prep column (Waters). Oligonucleotides were desalted using ethanol precipitation and quantification performed measuring the absorbance at 260 nm. Quality control of the synthesized oligonucleotides was performed by LC-MS.

### Antisense oligonucleotide treatment

SH-SY5Y cells with stable integration of a shRNA cassette for *TARDBP* were treated with 25 ng/ml doxycycline for 5 days and then transfected while replating into 12 well plates using Lipofectamine RNAiMax (6 μl per well) and ASOs at 50 nM final concentration. Following transfection doxycycline concentration was increased to 1 μg/ml and cells were harvested 5 days post-transfection. To test 18 nt MOE based ASOs, sh*TARDBP* SH-SY5Y cells were treated with 25 ng/mL doxycycline for 5 days, then 100 ng/mL doxycycline for 24 hours, plated into 96-well plates and transfected using Lipofectamine RNAiMax (0.72 μL per well) and ASOs at 10 nM or 50 nM final concentration. Cells were harvested 48 hours post-transfection.

SK-N-BE(2) cells with stable integration of a shRNA cassette for *TARDBP* were treated with 25 ng/ml doxycycline for 3 days and then transfected while replating into 12 well plates using Lipofectamine RNAiMax (6 μl per well) and ASOs (4 μl of 100 μM stock). Following transfection doxycycline concentration was increased to 1 μg/ml and cells were harvested 3 days post-transfection.

HaloTDP iNeurons were cultured for 2 weeks post-replating in neuronal media and then treated with 30 nM HaloPROTAC-E. ASOs (1 μM) were added directly to neuronal media to enable free uptake. ASO treatments were started at the same time as PROTAC treatment and were included in each media change for two weeks until cells were harvested.

### *UNC13A* exon 20-21 cryptic exon deletion

sgRNAs were designed to target 45 base pairs upstream of the acceptor splice site of the long cryptic exon (GGAAAAGTAAAAGATGTCTGAAT) and 10 base pairs downstream of the donor splice site (GAGATGGGTGAGTACATGGA). Sense and antisense bridging oligonucleotides `5’-AAAGTCCAGATTCAGGGTCACCTCCTCTGGGAAGCCCACCTTGGCCTCCBGGATGGATAGATGG ATGAGTTGGTGGGTAGATTCGTGGCTAGATGGATGA`-`3’` and 5’-TCATCCATCTAGCCACGAATCTACCCACCAACTCATCCATCTATCCATCDTGGAGGCCAAGGTGGGCTTCCCAGAGGAGGTGACCCTGAATCTGGACTTT-3’ which include degenerate base B (C,G, or T) and D (A,G, of T) were designed as homology-directed repair templates. WTC11 iPSC line (GM25256) and SH-SY5H were electroporated with 250 pmol of each Cas9 sgRNA (IDT) along with 20 μg recombinant Cas9 (IDT) and 1 pmol of each bridging oligonucleotide using the P3 Primary Cell 4-D-Nucleofector kit (Lonza) using the CA-137 program^34^. Following electroporation cells were maintained in media with 1x Alt-R HDR Enhancer V2 (IDT) for 24 hours.

### HaloTag editing of *TARDBP*

iPSCs were electroporated with 10 μg HaloTag-*TARDBP* homology-directed repair template, 500 pmol Cas12 crRNA (GGAAAAGTAAAAGATGTCTGAAT, IDT) and 20 μg recombinant Cas12 (IDT) using the P3 Primary Cell 4-D-Nucleofector kit (Lonza) and 4D-Nucleofector X unit (Lonza) using the CA-137 program. After electroporation iPSCs were plated in E8 Flex Media supplemented with 1x RevitaCell (Thermo) and 1x Alt-R HDR Enhancer V2 (IDT) for 24 hours and then expanded in E8 Flex Media. Cells were labelled with HaloTag-TMR ligand (Promega) and positive clones were selected for genotyping by PCR.

### Lentivirus production

sgRNA pU6-sgRNA EF1Alpha-puro-T2A-BFP vector (addgene 60955) was used to express the following sgRNAs:

sgCTRL-GTCCACCCTTATCTAGGCTA

sg*TARDBP*-GGGAAGTCAGCCGTGAGACC

sg*UNC13A*-GGAACCAAGATGGCCGGTGG

sgRNA transfer plasmid with packaging (pCMVΔ8.91) and VSVG envelope (pMD.G) plasmids were transfected into HEK293T cells (Horizon Discovery) with TransIT-293 (Mirus). Media was collected on day 2 and 3 post-transfection, combined, and concentrated 1:10 with Lenti-X concentrator (Takara Bio).

### RT-qPCR

RNA was extracted from SH-SY5Y cells, SK-N-BE(2) cells and iNeurons with an RNeasy kit (Qiagen) using the manufacturer’s protocol including homogenization of samples with QIAshredder spin columns (Qiagen). RNA concentrations were measured by Nanodrop and 500 ng of RNA was used for reverse transcription. First-strand cDNA synthesis was performed with RevertAid (Thermo K1622) using random hexamer primers and following the manufacturer’s protocol including all optional steps. Gene expression analysis was performed by RT-qPCR using Taqman Multiplex Universal Master Mix (Thermo 4461882) with custom IDT RT-qPCR assays (Sup Methods Table) and TaqMan assays (GAPDH-Jun assay 4485713) on a QuantStudio 5 Real-Time PCR system (Applied Biosystems) and quantified using the ΔΔCt method^35^. For RT-qPCR based on intercalating dyes PowerUp SYBR Green master mix (Thermo) was used with gene-specific primers (Sup Methods Table).

### RT-PCR

RNA extraction and cDNA synthesis were performed as in RT-qPCR methods. The *UNC13A* CE was amplified with a forward primer in exon 19 5’-CAGACGATCATTGAGGTGCG-3’and reverse primer in exon 22 5’-ATACTTGGAGGAGAGGCAGG-3’using Q5 High Fidelity Master Mix (NEB). PCR products were resolved on a TapeStation 4200 (Agilent) and bands were quantified with TapeStation Systems Software v3.2 (Agilent). For the primary screen of ASOs in SK-N-BE(2) cells, PCR products were analysed with a QIAxcel capillary electrophoresis machine. Raw data was exported and analyzed with the QIAxcelR package (v0.1) (https://github.com/Delayed-Gitification/QIAxcelR).

### Amplicon sequencing

Genomic DNA from SH-SY5Y cells and iPSCs was extracted with the Monarch Genomic DNA Purification Kit (NEB). PCR was performed with Q5 Hot Start High-Fidelity Master Mix (NEB) and purified with Monarch PCR cleanup kit (NEB). Purified amplicons were sequenced by Plasmidsaurus. Raw FASTQ reads were mapped to *UNC13A* and *TARDBP* genes using Minimap2 with the -x splice flag enabled^36^. Results were visualised with IGV browser.

### Western blot analysis

Cells were harvested with RIPA buffer (Thermo) with cOmplete mini protease inhibitors (Sigma) and protein concentration was quantified using the BCA assay (Pierce). Protein lysates were resolved on 4-12 % precast Bis-Tris gels (Thermo) or poured 7% Bis-Tris gels and transferred to 0.2 μm PVDF membranes (Millipore). Membranes were blocked with 5% non-fat milk in TBST and then probed overnight with primary antibodies in 5% non-fat milk in TBST at 4°C. Blots were washed 3 times with TBST and subsequently probed with HRP-conjugated secondary antibodies at room temperature for 1 hour. Goat anti-Mouse HRP (Bio-Rad 1706516) 1:10,000 Goat anti-Rabbit HRP (Bio-Rad 1706515) and developed with Chemiluminescent substrate (Merck Millipore WBKLS0500) on a ChemiDoc Imaging System (Bio-Rad). Band intensity was measured with ImageJ (NIH version 2.0.0-rc-69).

### Multi-electrode array recordings

Maestro 96-well cytoview multi-electrode array plates (Axion Biosystems M768-tMEA-96B) were dropwise coated in 0.05 mg/ml poly-D-lysine (Gibco A38904-01) at RT for 2 hours, washed 3x in 50 μl ddH_2_O and left to dry overnight. Wells were then dropwise coated with 10 ng/ml laminin for 1-2 hours at 37°C. 50,000 neurons and 12,500 primary rat astrocytes were dropwise plated in synaptic media containing ROCK inhibitor (Tocris) at 1 nM. Cells were left to attach for 1.5 hours before media containing CultureONE supplement (Gibco) was added for 3 days. Half media changes (without ROCK inhibitor or CultureONE) were thereafter performed every 2-3 days and HaloPROTAC-E 30-300 nM was added to cultures with each media change starting on DIV 14. MEA plates underwent recordings once every 2-3 days on the Maestro Pro (Axion Biosystems). Plates were left inside the Maestro Pro for 15 minutes preceding each recording to allow for acclimatisation and recordings occurred immediately after and continued for 15 minutes. Data was processed using the Neural Metric Tool (Axion Biosystems). Active electrode criteria were set to 5 spikes/min. Single electrode burst parameters used an ISI threshold, consisting of a minimum of 5 spikes and a max ISI of 100 ms. Network bursts used envelope analysis, with a threshold factor of 1.25, min IBI of 100 ms, 35% min electrode and a burst exclusion of 75%. All synchrony metrics were measured with a synchrony window of 20 ms. Wells with fewer than 3 active electrodes were excluded from the analysis. Data was normalised to the maximum mean of each control condition and statistical analysis was performed on area under the curve values for time-course data between day 28 and the last day of neuronal survival.

### Electrophysiology recordings

For whole-cell patch clamp recordings, 500k i3 neurons were plated onto 18 mm No.1 coverslips (Deckglaser) coated with 50 μg/mL poly-D-lysine (Gibco) and mouse 10 μg/mL laminin (Gibco). Primary astrocytes obtained from E18 rat cortices via 15 min trypsinisation (10 mg/ml; Gibco), were plated 1 week prior and treated with culture ONE supplement (Gibco) 2 days prior. Miniature excitatory postsynaptic currents (mEPSCs) were recorded at 4-weeks maturation on rat astrocytes, in extracellular solution containing: 136 mM NaCl, 2.5 mM KCl, 10 mM HEPES, 1.3 mM MgCl_2_, 2 mM CaCl_2_, 10 mM Glucose, pH 7.3, 300 mOsm, supplemented with 10 μM tetrodotoxin (TTX) and 10 μM gabazine (GZ). Patching pipettes were filled with cesium methanosulfate-based internal solution: 135 mM CsMeSO_3_, 10 mM HEPES, 10 mM Na_2_-Phosphocreatine, 5 mM Glutathione, 4 mM MgCl_2_, 4 mM Na_2_ATP, 0.4 mM NaGTP supplemented with 5 μM QX-314. Pipettes were pulled from borosilicate glass (O.D. 1.5 mm, I.D. 0.86 mm, Sutter instruments) to a resistance of between 3-5 MΩ. Whole-cell patch clamp recordings were then made without whole-cell compensation applied, for 5 minutes at the soma. Cells were held at -70 mV using a Multiclamp 700B amplifier (Molecular Devices) and the data were acquired using a Digidata 1440A digitizer (Molecular Devices). All recordings were carried out on a heated stage set to 37°C. Data was acquired with Clampex software (Molecular Devices) and Axon Multiclamp Commander Software (Molecular Devices), data was sampled at a rate of 20 kHz and filtered at 10 kHz. mEPSCs were analysed in a fully automated manner using Clampfit 10.7 software (Molecular devices). Passive membrane properties were measured by applying a subthreshold voltage step and analysed using custom MATLAB scripts. Series resistance (Rs) was calculated as proportional to the current response amplitude, membrane capacitance (Cm) was estimated from the transient time constant by fitting the current transients to exponential functions and membrane resistance (Rm) was estimated from the steady-state response following the current transients. Passive membrane properties and series resistance were measured at the beginning and at the end of all recordings to ensure proper access (RS >30 mΩ) had been maintained.

### TDP-43 immunofluorescence

20k neurons were grown poly-D-lysine/laminin-coated coverslips. Neurons were fixed at DIV 28 in 4% PFA, 4% sucrose for 15 minutes. Cells were then permeabilised in 0.1% Triton X-100 in PBS for 10 minutes. Primary antibodies, mouse anti-TDP-43 (Abcam ab104223) and chicken anti MAP2 (Abcam ab5392) were diluted in PBS at 1:1,000 and 1:10,000 respectively and incubated overnight at 4°C. Cells were washed 3x in PBS. Secondary antibodies, goat anti-mouse IgG (H+L) AlexaFluor 488 (ThermoFisher) and goat anti-chicken IgY (H+L) Alexafluor 647 (ThermoFisher) were diluted in PBS 1:1000 and incubated at RT for 1hr. Cells were washed 3x in PBS, labeled with DAPI (1:1,000) for 5 minutes at RT, and mounted in ProLong Gold mounting medium (ThermoFisher). Cells were imaged using a Zeiss 980 Airyscan confocal at 63x.

### UNC13A Immunofluorescence

500k neurons were grown on 50k primary rat astrocytes on poly-D-lysine/laminin-coated coverslips. Neurons were fixed at DIV 35 in 4% PFA for 10 minutes. Cells were then permeabilised in 0.1% Triton X-100 in PBS for 10 minutes. Primary antibodies, guinea pig anti-Munc13-1 (Synaptic Systems - 126-104) and mouse anti Synapsin 1 (Synaptic Systems - 106-011) were diluted in PBS at 1:500 and 1:1,000 respectively and incubated for 1hr at RT. Cells were washed 3x in PBS. Secondary antibodies, goat anti-mouse IgG (H+L) AlexaFluor 488 (ThermoFisher) and goat anti-guinea-pig IgG (H+L) Alexafluor 647 (ThermoFisher) were diluted in PBS 1:1000 and incubated at RT for 1hr. Cells were washed 3x in PBS and mounted in Mowiol-4-88 (PolySciences) mounting medium containing 1:1,000 DAPI (ThermoFisher). Cells were imaged using a Zeiss 980 Airyscan confocal at 63x with a 3x crop. Z-stacks of 7x 0.5 μm intervals were taken. FIJI was used to analyse mean UNC13A intensity based on manually selected synapsin-positive ROIs.

### Statistical Analysis

Datasets were assessed for normality. For normally distributed data, t-tests were used for statistical comparison of two groups. One-way ANOVA tests with post-hoc multiple pairwise comparison of means was used for statistical analysis of three or more groups. For non-normally distributed data, nonparametric tests were used. Statistical tests were carried out using Prism (Graphpad) or with the R statistical library. Sample size and *p* values are specified in figure legends.

### Materials availability

The sh*TARDBP* SH-SY5Y cell line, sh*TARDBP* SK-N-BE(2) cell line, *UNC13A* cryptic exon deletion SH-SY5Y cell line, *UNC13A* cryptic exon deletion iPSC line, and Halo-TDP-43 iPSC line are available from PF under a material transfer agreement with UCL.

### Data availability

Source data is provided with this paper.

### Code availability

MATLAB codes for electrophysiology are available from JB.

## Acknowledgments

This work was supported by UK Medical Research Council Senior Clinical Fellowship MR/M008606/1 (PF); Motor Neurone Disease Association Lady Edith Wolfson Fellowship MR/S006508/1 (PF); National Institute of Health U54NS123743 (PF); The Sigrid Rausing Neurogenetic Therapies Programme (PF, MJK, OGW); LifeARC (PF); Wellcome Trust Clinical Training Fellowship 102186/B/13/Z (PRM); This research was funded in whole, or in part, by the Wellcome Trust (215508/Z/19/Z) to JB. For the purpose of open access, the author has applied a CC BY public copyright licence to any Author Accepted Manuscript version arising from this submission; Biotechnology and Biological Sciences Research Council project grant BB/S000526/1 (JB); NATA is supported by the Medical Research Council (MC_PC_20061) (PLO)

## Author contributions

Conceptualization: PF, JB, MJK, PH, OGW, GS

Methodology: MJK, PH, ER, REJ, MZ, OGW, JEP, AFCS

Investigation: MJK, PH, ER, REJ, MZ, OGW, SB, PRM, JEP, AS, AFCS

Visualization: MJK, PH, ER, OGW

Funding acquisition: PF, JB, PLO, EMCF, GS, MS

Supervision: PF, JB, GS, EMCF, PLO, MS

Writing – original draft: MJK, PH, PF, JB

Writing – review & editing: ER, REJ, MZ, OGW, SB, PRM, JEP, AS, AFCS, PLO, EMCF, GS, MS

