## Supplemental Figures and Tables for "Loss of TDP-43 induces synaptic dysfunction that is rescued by *UNC13A* splice-switching ASOs"

### Extended Data Figures and Table

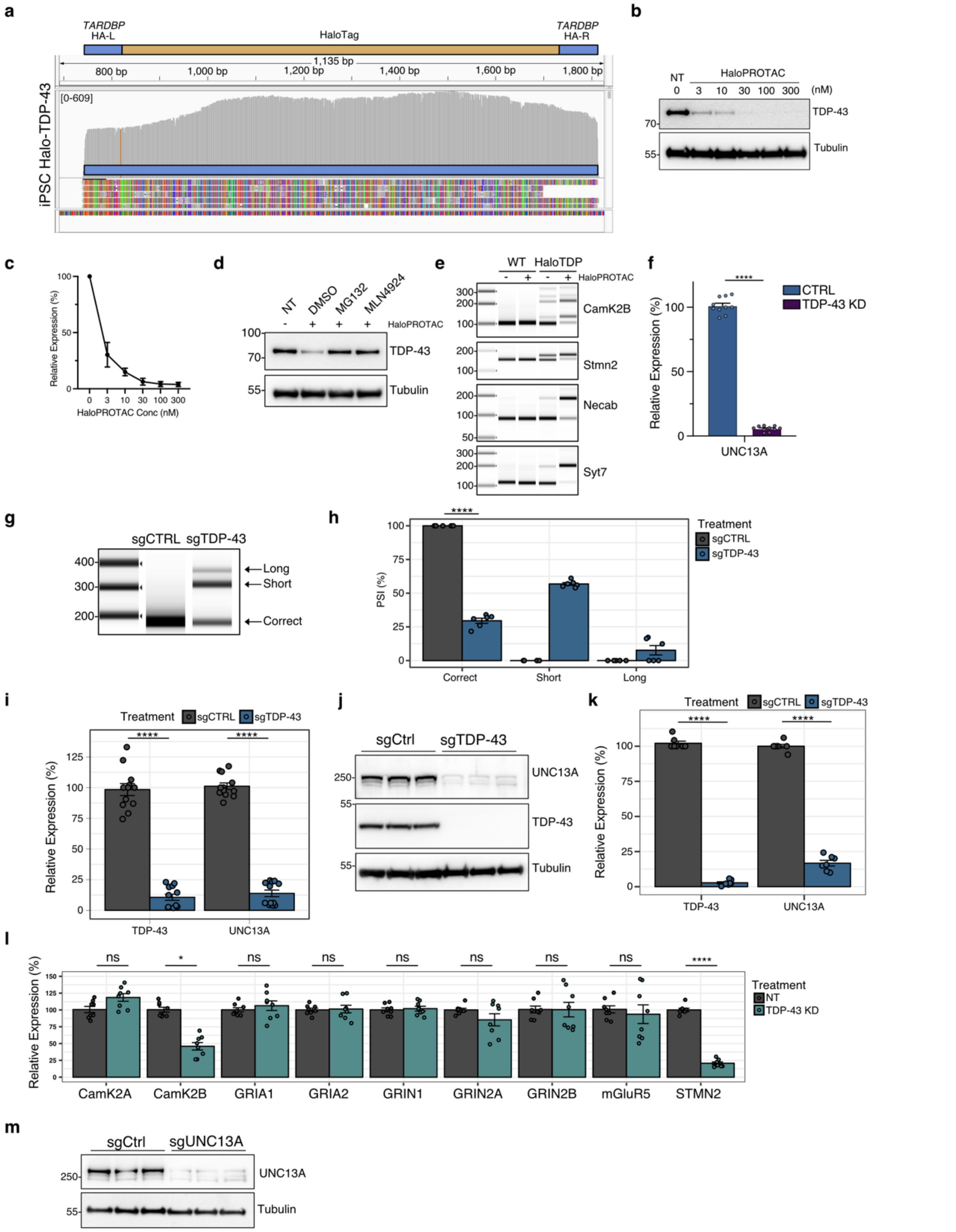


**Extended Data Figure 1 | CRISPRi mediated knockdown of TDP-43 in iNeurons. a**, Amplicon sequencing of PCR products from Fig. 1A indicates correct HaloTag editing of *TARDBP*. **b**, HaloTDP iNeurons were treated with the indicated concentration of HaloProtac for 4 days and processed for western blot analysis indicating a dose response curve of HaloProtac on TDP-43 knockdown. **c**, Quantification of western blots in (**b**) *n*=3 biological replicates. **d**, HaloTDP iNeurons were treated with 30 nM HaloProtac for 3 hours in the presence of DMSO, proteasome inhibitor MG132, or NEDD8 E1 inhibitor MLN4924. **e**, RT-PCR analysis of WT and Halo-iNeurons shows the Halo tag induces a mild loss of TDP-43 function and can induce cryptic exon inclusion in the indicated transcripts. **f**, RT-qPCR analysis of correctly spliced *UNC13A* at exon 20-21 junction. *n*=10 biological replicates from 3 experiments. **g-k**, Analysis of control and CRISPRi-mediated TDP-43 knockdown in DIV 28 iNeurons. (**g**) RT-PCR analysis of *UNC13A* splicing at exon junction 20-21 shows the inclusion of a long and short cryptic exon after TDP-43 KD. (**h**) Quantification of results in (**g**) *n*=6 from 3 experiments. (**i**) RT-qPCR analysis shows a reduction of correctly spliced *UNC13A* at exon 20-21 junction following TDP-43 KD. (**j**) Western blot analysis indicates a dramatic decrease in UNC13A protein after TDP-43 KD. (**k**) Quantification of results in (**j**) *n*=7 from 3 experiments. **l**, RT-qPCR analysis of DIV 38 HaloTDP iNeuron cultures indicates similar expression of glutamate receptors. A cryptic exon in *CamK2B* and cryptic polyadenylation in *STMN2* results in reduction of mRNA transcripts. *n*=8 from 2 experiments. **m**, Western blot analysis of control and CRISPRi-mediated UNC13A knockdown in iNeurons. Graphs in (**f**) (**h**) (**i**) (**k**) and (**l**) represent mean ± s.e.m. Statistics are two-sided Student's *t* test with additional Bonferroni adjustment for multiple comparisons in (**l**) *$P$ < 0.05; ** $P$ <0.01; *** $P$ <0.001; **** $P$ < 0.0001; ns (not significant).

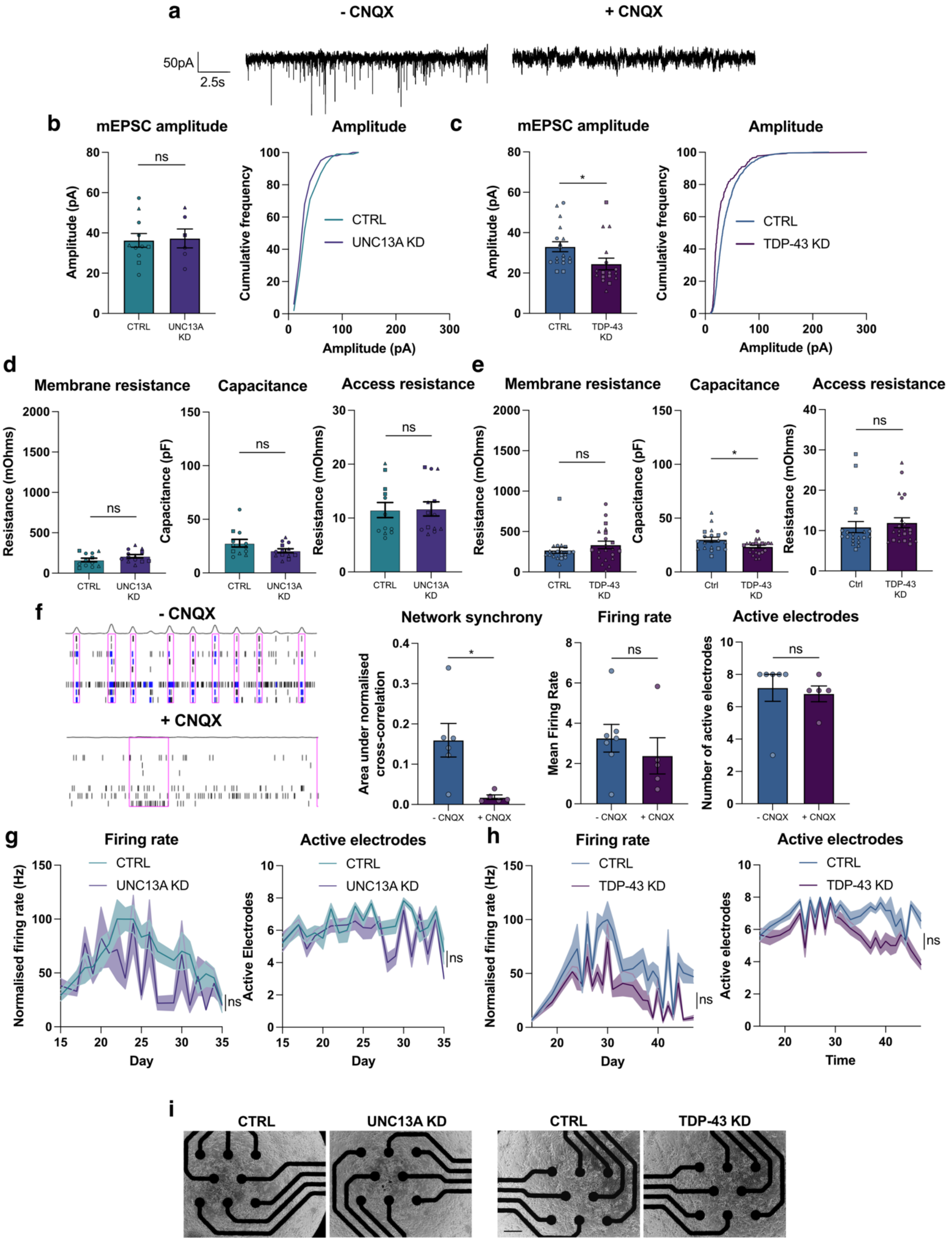


**Extended Data Figure 2 | Additional electrophysiological parameters following UNC13A and TDP-43 knockdown. a**, AMPAR blockade using CNQX abolishes mEPSC activity. **b**, mEPSC amplitude from control (*n*=12) and UNC13A depleted (*n*=13) iNeurons pooled from 3 experiments. **c**, mEPSC amplitude from control (n=20) and TDP-43 knockdown (n=22) HaloTDP iNeurons from 3 experiments. **d**, Quantification of passive membrane properties for UNC13A knockdown iNeurons. **e**, Quantification of passive membrane properties for TDP-43 knockdown iNeurons. **f**, AMPAR blockade using CNQX abolishes network synchrony on multielectrode array recordings, with minimal effects on firing rate and number of active electrodes. **g**, Quantification of firing rate and number of active electrodes following UNC13A knockdown. **h**, Quantification of firing rate and number of active electrodes following TDP-43 knockdown. **i**, Phase contrast images showing cell coverage on multielectrode array plates. Scale bar = 350 µm. Graphs for (**b**) (**c**) (**d**) (**e**) (**f**) represent mean ± s.e.m. Statistics for (**b**) (**c**) (**d**) and (**f**) are two-sided Student's *t* tests. Statistics for (**g**) and (**h**) are paired *t* tests. $*P < 0.05$; ns (not significant).

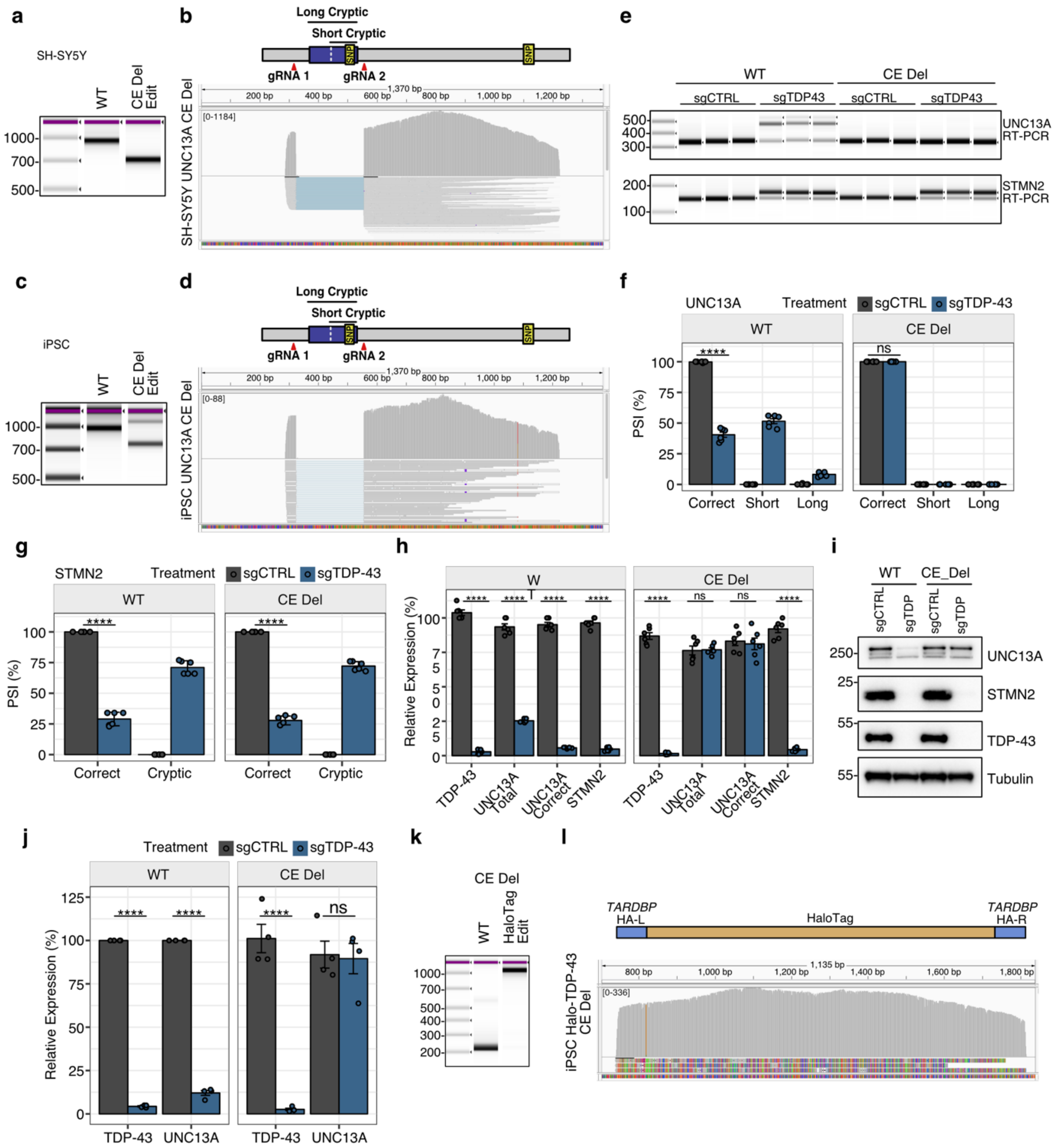


**Extended Data Figure 3 | *UNC13A* cryptic exon deletion rescues *UNC13A* splicing/expression following TDP-43 knockdown. a**, PCR from genomic DNA of WT SH-SY5Y cells and CE Del SH-SY5Y cells indicated successful deletion of the genomic region of *UNC13A* containing the sequence corresponding to the exon 20-21 cryptic exon. **b**, Amplicon sequencing of PCR products in (**a**). **c**, PCR from genomic DNA of WT iPSCs and CE Del iPSCs indicated successful deletion of the genomic region of *UNC13A* containing the sequence corresponding to the exon 20-21 cryptic exon. **d**, Amplicon sequencing of PCR products in (**c**) **e-j**, Analysis of WT and CE Del iNeurons following CRISPRi knockdown with CTRL or TDP-43 sgRNAs. (**e**) RT-PCR analysis shows *UNC13A* cryptic deletion rescues *UNC13A* exon 20-21 splicing following TDP-43 knockdown, but not cryptic polyadenylation of *STMN2* (**f**). Quantification of *UNC13A* splicing in figure (**e**) *n*=6 biological replicates from 2 experiments. (**g**) Quantification of *STMN2* splicing in figure (**e**) *n*=6 biological replicates from 2 experiments. (**h**) RT-qPCR analysis shows *UNC13A* CE Del prevents the loss of *UNC13A* transcripts but not *STMN2* transcripts in iNeurons after TDP-43 knockdown. *UNC13A* correct RT-qPCR assay detects correct splicing at *UNC13A* exon 20-21 junction. *n* = 6 biological replicates from 2 experiments. (**i**) Western blot analysis shows *UNC13A* CE Del rescues the loss of UNC13A protein following TDP-43 knockdown. (**j**) Quantification of western blots in (**i**) *n*=4 biological replicates from 2 experiments. **k**, PCR of genomic DNA of *UNC13A* CE Del iPSCs with primers flanking exon 1 of *TARDBP* indicates successful edit of *TARDBP* with HaloTag. **l**, Amplicon sequencing of HaloTag PCR product in (**k**). Graphs for (**f**) (**g**) (**h**) and (**j**) represent mean ± s.e.m. Statistics are One-way ANOVA with Tukey multiple comparison test. *$P < 0.05$; ** $P < 0.01$; *** $P < 0.001$; **** $P < 0.0001$; ns (not significant).

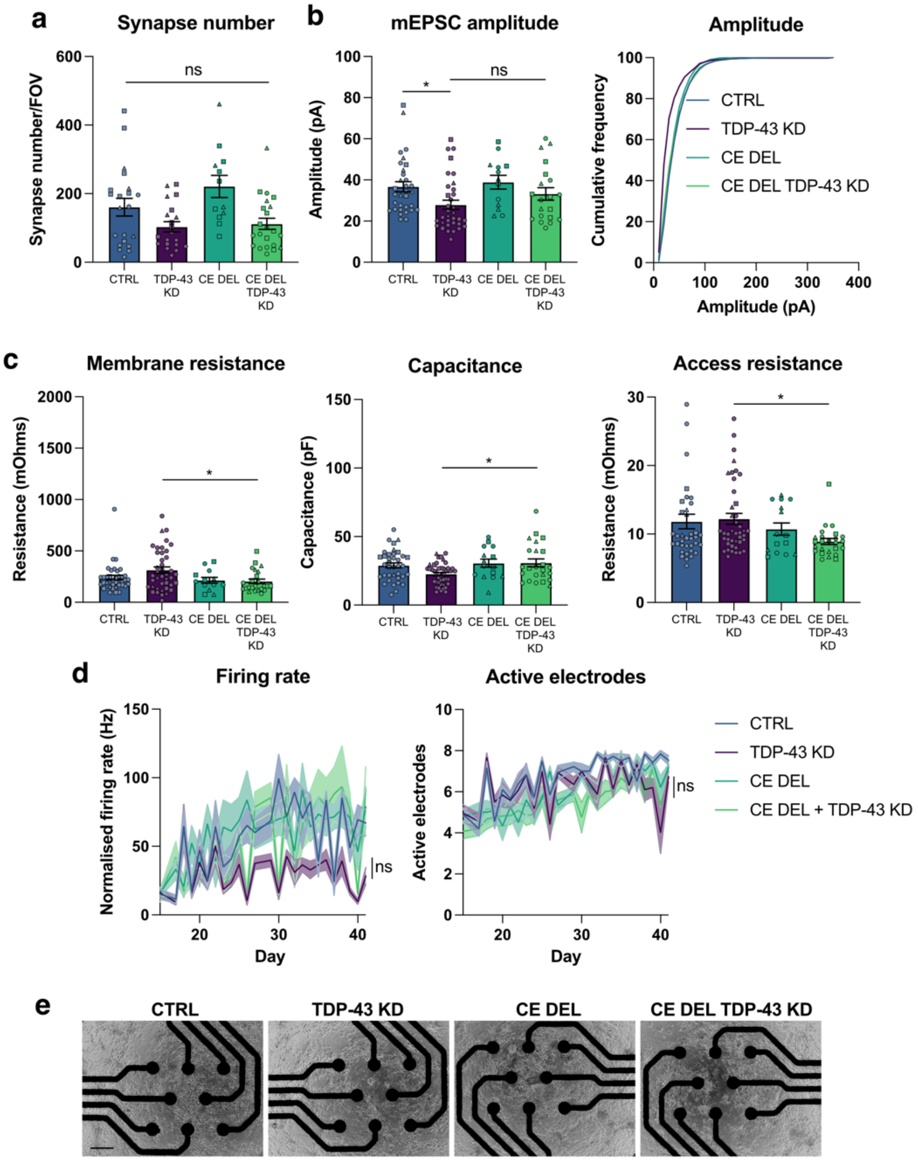


**Extended Data Figure 4 | Additional synaptic and electrophysiological parameters following genomic deletion of *UNC13A* CE. a,** Immunofluorescence quantification of synapse number and size based on synapsin labelling in control *n*=21, TDP-43 KD *n*= 20, CE Del *n*=12 and CE Del TDP-43 KD *n*=22 fields of view from 3 experiments. **b,** mEPSC amplitude from control *n*=17, TDP-43 knockdown *n*=22, CE Del *n*=15, CE Del TDP-43 knockdown *n*=28 iNeurons pooled from 3 experiments. **c,** Quantification of passive membrane properties. **d,** Quantification of multielectrode array mean firing rates and number of active electrodes for control, TDP-43 KD, CE Del, CE Del TDP-43 KD conditions *n*=18 wells from 3 experiments. **e,** Phase contrast images showing cell coverage on multielectrode array plates. Scale bar = 350 µm. Graphs for (**a**) (**b**) (**c**) (**d**) represent mean ± s.e.m. Statistics are One-way ANOVA with Dunnet's multiple comparison test. *$P < 0.05$; ns (not significant).

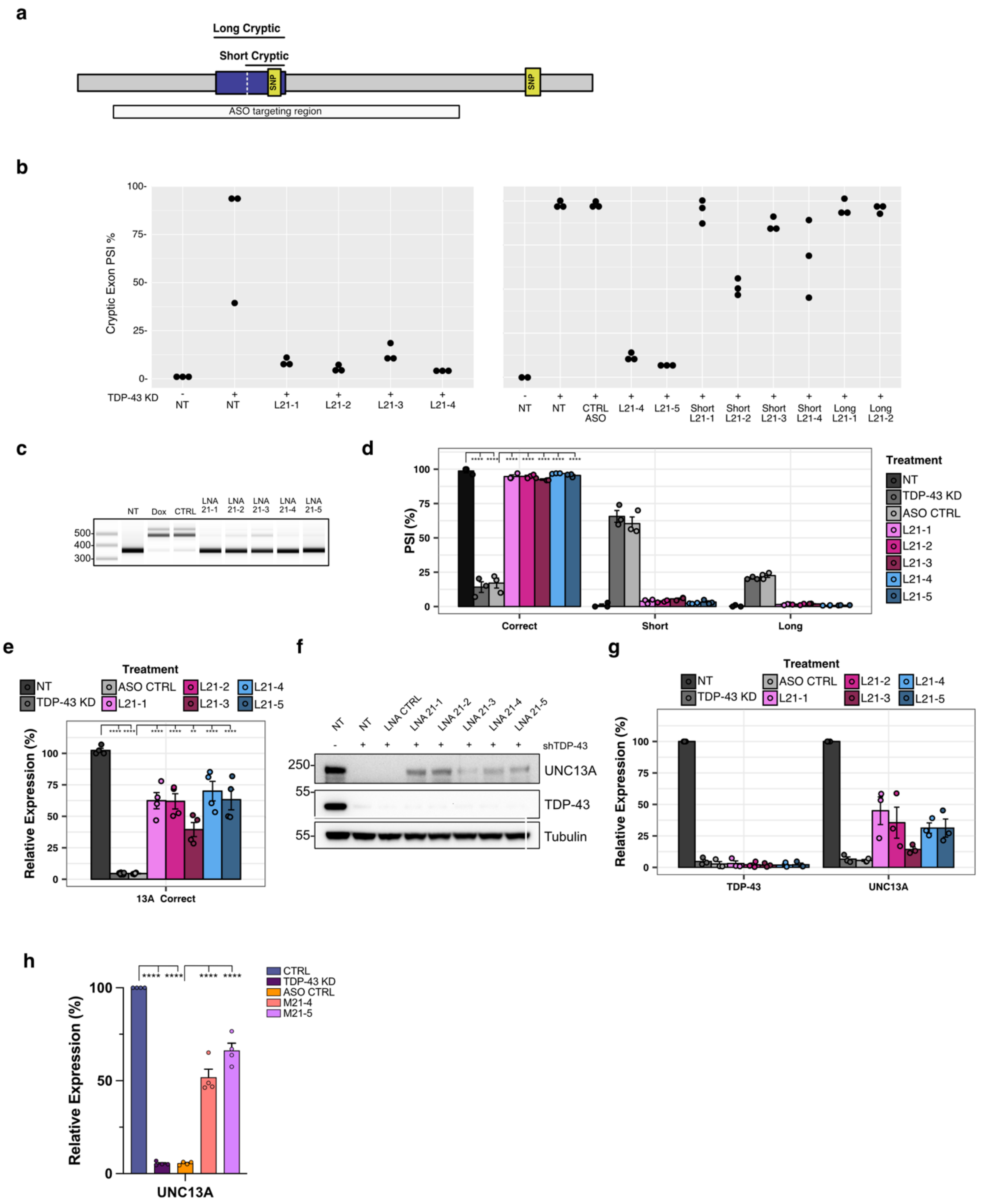


**Extended Data Figure 5 | ASOs targeting the *UNC13A* cryptic exon rescues *UNC13A* splicing/expression following TDP-43 knockdown.** **a**, Schematic of *UNC13A* cryptic exon. **b**, Quantification of RT-PCR products following treatment of SK-N-BE(2) cells with indicated ASOs. **c**, RT-PCR analysis of *UNC13A* splicing in SH-SY5Y cells shows ASOs prevent cryptic splicing after TDP-43 knockdown. **d**, Quantification of results in (C) *n*=3 biological replicates from 2 experiments. **e**, RT-qPCR analysis of *UNC13A* shows an ASO mediated rescue of *UNC13A* RNA after TDP-43 KD in SH-SY5Y cells. *n*=4 biological replicates from 2 experiments. **f**, Western blot analysis shows rescue of UNC13A protein following ASO treatment after TDP-43 KD in SH-SY5Y cells. **g**, Quantification of blots from (**f**). *n*=3 biological replicates from 3 experiments. **h,** RT-qPCR analysis of *UNC13A* shows an ASO mediated rescue of *UNC13A* RNA after TDP-43 KD in HaloTDP iNeurons. *n*=4 biological replicates from 2 experiments. Graphs for (**d**) (**e**) (**g**) and (**h**) represent mean ± s.e.m. Statistics are One-way ANOVA with Tukey multiple comparison test. **P* < 0.05; ** *P* <0.01; *** *P* <0.001; **** *P* < 0.0001; ns (not significant).

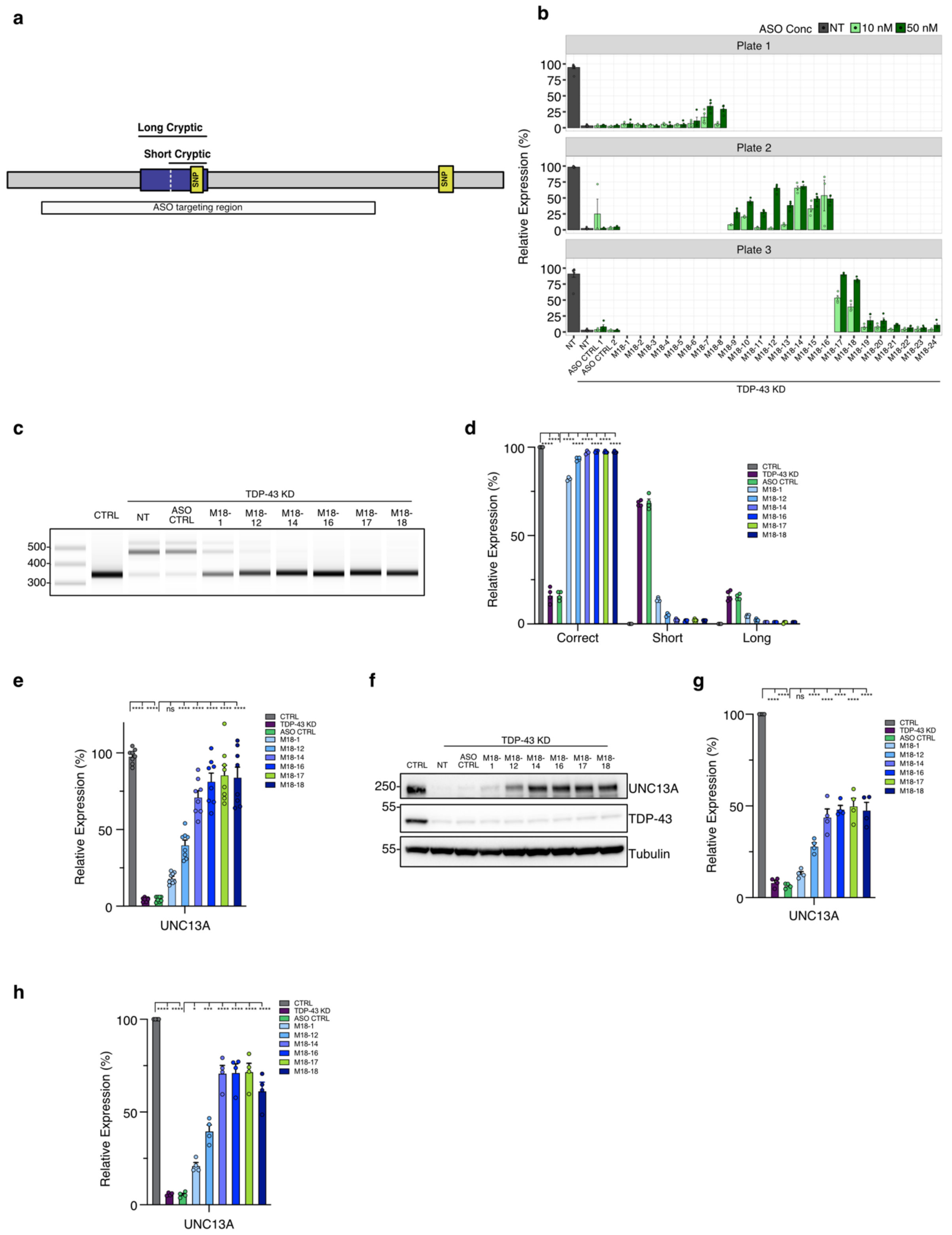


**Extended Data Figure 6 | Improved ASOs targeting the *UNC13A* cryptic exon rescues *UNC13A* splicing/expression following TDP-43 knockdown.** **a**, Schematic of *UNC13A* cryptic exon. **b**, Quantification of RT-PCR products following treatment of SH-SY5Y cells with indicated ASOs. **c**, RT-PCR analysis of *UNC13A* splicing in SH-SY5Y cells shows ASOs prevent cryptic splicing after TDP-43 knockdown. **d**, Quantification of results in (**c**) *n*=3 biological replicates from 2 experiments. **e**, RT-qPCR analysis of *UNC13A* shows an ASO mediated rescue of *UNC13A* RNA after TDP-43 KD in SH-SY5Y cells. *n*=4 biological replicates from 2 experiments. **f**, Western blot analysis shows rescue of UNC13A protein following ASO treatment after TDP-43 KD in SH-SY5Y cells. **g**, Quantification of blots from (**f**). *n*=3 biological replicates from 3 experiments. **h**, RT-qPCR analysis of *UNC13A* shows an ASO mediated rescue of *UNC13A* RNA after TDP-43 KD in Halo-iNeurons. *n*=4 biological replicates from 2 experiments. Graphs for (**d**) (**e**) (**g**) and (**h**) represent mean ± s.e.m. Statistics are One-way ANOVA with Tukey multiple comparison test. **P* < 0.05; ** *P* <0.01; *** *P* <0.001; **** *P* < 0.0001; ns (not significant).

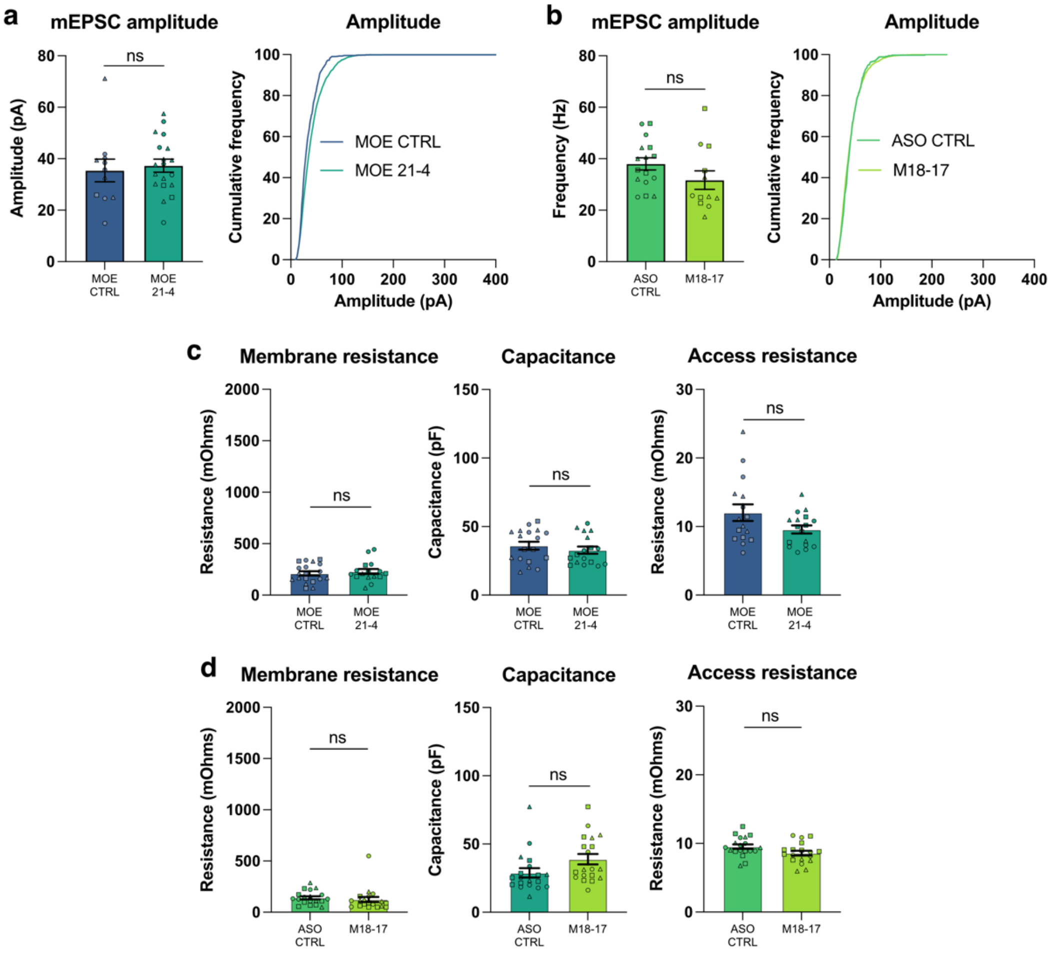


**Extended Data Figure 7 | Additional electrophysiological parameters following ASO treatments.** **a**, Quantification of mEPSC amplitude in TDP-43 KD iNeurons treated with MOE CTRL ASO (*n*=17) and MOE 21-4 ASO (*n*=18) from 3 experiments. **b,** Quantification of mEPSC amplitude in TDP-43 KD iNeurons treated with CTRL ASO (*n*=19) and M18-17 ASO (*n*=19) from 3 experiments. **c,** Passive membrane properties for TDP-43 KD iNeurons treated with MOE CTRL ASO (*n*=17) and MOE 21-4 ASO (*n*=18) from 3 experiments. **d,** Passive membrane properties in TDP-43 KD iNeurons treated with CTRL ASO (*n*=19) and M18-17 ASO (*n*=19) from 3 experiments. Graphs for (**a**) (**b**) (**c**) and (**d**) represent mean ± s.e.m. Statistics are two-sided Student's *t* test. ns (not significant).

**Extended Data Table 1 | Key Reagents**

| Reagent | Supplier | Catalog # |
|---|---|---|
| XAV939 | Cambridge Bioscience | SM38-10 |
| LDN-193189 hydrochloride | Cambridge Bioscience | 19396-5mg-CAY |
| SB431542 | Cambridge Bioscience | SM33-50 |
| Recombinant Cas9 | IDT | Alt-R® S.p. Cas9 Nuclease V3 |
| Recombinant Cas12 | IDT | Alt-R® A.s. Cas12a (Cpf1) Ultra |
| HDR Enhancer V2 | IDT | NA |
| Trans-IT 293 | Mirius | MIR2700 |
| dibutyryl cAMP | Merck Sigma | D0627-100MG |
| Doxycycline Hyclate | Merck Sigma | D9891 |
| L-Ascorbic Acid | Merck Sigma | A0278-25G |
| Poly-D-lysine hydrobromide >300,000 MW | Merck Sigma | P7405 |
| Poly-L-ornithine hydrobromide 30,000-70,000 MW | Merck Sigma | P3655 |
| Puromycin | Merck Sigma | P8833 |
| BDNF | PeproTech | 450-02 |
| GDNF | PeproTech | 450-10 |
| Accutase | ThermoFisher | A1110501 |
| cultureOne | ThermoFisher | A3320201 |
| Geltrex | ThermoFisher | A14133-01 |
| Laminin | ThermoFisher | 23017015 |
| RevitaCell | ThermoFisher | A2644501 |
| TrypLE | ThermoFisher | 12605010 |
| Versene | ThermoFisher | 15040066 |
| ROCK inhibitor (Y-27632) | Tocris | 1245/10 |
| Halo-Protac E | University of Dundee DSTT | HALO-PROTAC-E active |
| Tetrodotoxin | Tocris | 1069 |
| Gabazine | Tocris | 1262 |

**Extended Data Table 2 | DNA Oligonucleotides sequences**

| Name | Sequence 5'-3' | Use | Supplier | Catalog Number |
|---|---|---|---|---|
| UNC13A_For | CAGACGATCATTGAGGTGCG | RT-PCR | IDT | NA |
| UNC13A_Rev | ATACTTGGAGGAGAGGCAGG | RT-PCR | IDT | NA |
| STMN2_For | GCTCTCTCCGCTGCTGTAG | RT-PCR | IDT | NA |
| STMN2_Rev | CGAGGTTCCGGGTAAAAGCA | RT-PCR | IDT | NA |
| STMN2_Cryptic_Rev | CTGTCTCTCTCTCTCGCACA | RT-PCR | IDT | NA |
| UNC13A_Total_For | TGATGTTGACCTCGATGAACG | RT-qPCR | IDT | Hs.PT.58.1883136 |
| UNC13A_Total_Rev | TCTGTCCATGTTGAGCTGTTC | RT-qPCR | IDT | Hs.PT.58.1883136 |
| UNC13A_Total_Probe | /56-FAM/AGCCACCAC/ZEN/TTTCACTGTGACCTT/3IABkFQ/ | RT-qPCR | IDT | Hs.PT.58.1883136 |
| UNC13A_Correct_For | GGACAAGCGAACTGACAAATC | RT-qPCR | IDT | NA |
| UNC13A_Correct_Rev | ACAGGTTCTCATGCAGACAG | RT-qPCR | IDT | NA |
| UNC13A_Correct_Probe | /5SUN/ATCAAAGGC/ZEN/GAGGAGAAGGTGGC/3IABkFQ/ | RT-qPCR | IDT | NA |
| Gapdh-Jun | NA | RT-qPCR | Thermo | 4485713 |
| CamK2A_For | TCAATCAGCTGCTCTGTCAC | RT-qPCR | IDT | Hs.PT.56a.28027747 |
| CamK2A_Rev | CCAGTTCCAGCGTTCAGTT | RT-qPCR | IDT | Hs.PT.56a.28027747 |
| CamK2B_For | TCTTGGCTGCATACTCATGG | RT-qPCR | IDT | Hs.PT.56a.20211778.g |
| CamK2B_Rev | CTTCACCGACGAGTACCAG | RT-qPCR | IDT | Hs.PT.56a.20211778.g |
| GRIA1_For | CTTAATCGAGTTCTGCTACAAATCC | RT-qPCR | IDT | Hs.PT.58.15517507 |
| GRIA1_Rev | GTATGGCTTCGTTGATGGTTG | RT-qPCR | IDT | Hs.PT.58.15517507 |
| GRIA2_For | GGTACGACAAAGGAGAGTGC | RT-qPCR | IDT | Hs.PT.58.25075751 |
| GRIA2_Rev | CCCGACAAGGATGTAGAATACTC | RT-qPCR | IDT | Hs.PT.58.25075751 |
| GRIN1_For | CTCCTGGAAGATTCAGCTCAA | RT-qPCR | IDT | Hs.PT.58.39141804 |
| GRIN1_Rev | GTGGATGGCTAACTAGGATGG | RT-qPCR | IDT | Hs.PT.58.39141804 |
| GRIN2A_For | CAAGAAGTAATGGCACCGTCT | RT-qPCR | IDT | Hs.PT.58.26949410 |
| GRIN2A_Rev | GCAGAAACAATGAGCAGCATC | RT-qPCR | IDT | Hs.PT.58.26949410 |
| GRIN2B_For | CTTCATAGAGACAGGCATCAGT | RT-qPCR | IDT | Hs.PT.58.40419546 |
| GRIN2B_Rev | CATCACAAACATCATCACCCATAC | RT-qPCR | IDT | Hs.PT.58.40419546 |
| mGluR5_For | TGTGAGAAAGGCCAGATCAAG | RT-qPCR | IDT | Hs.PT.58.40025787 |
| mGluR5_Rev | TGCCTTGCATGTGTACTCATC | RT-qPCR | IDT | Hs.PT.58.40025787 |
| STMN2_For | CCACGAACTTTAGCTTCTCCA | RT-qPCR | IDT | Hs.PT.58.5075784 |
| STMN2_Rev | GCCAATTGTTTCAGCACCTG | RT-qPCR | IDT | Hs.PT.58.5075784 |

**Extended Data Table 3 | Antibodies**

| Antibody | Supplier; Catalog Number | Use | Dilution |
|---|---|---|---|
| Rabbit anti UNC13A | Synaptic Systems; 126 103 | WB | 1:2,000 |
| Rabbit anti TDP-43 | ProteinTech; 10782-2-AP | WB | 1:2,000 |
| Mouse anti Tubulin | ProteinTech; 66031-1-Ig | WB | 1:2,000 |
| Rabbit anti STMN2 | ProteinTech; 10586-1-AP | WB | 1:2,000 |
| Goat anti Mouse IGG-HRP conjugate | BioRad; 1706516 | WB | 1:10,000 |
| Goat anti Rabbit IGG-HRP conjugate | BioRad; 1706515 | WB | 1:10,000 |
| Mouse anti TDP-43 | Abcam; ab104223 | IF | 1:1,000 |
| Mouse anti Synapsin | Synaptic Systems; 106 011 | IF | 1:1,000 |
| Guinea Pig anti UNC13A | Synaptic Systems; 126 104 | IF | 1:500 |
| Chicken anti MAP2 | Abcam; ab5392 | IF | 1:10,000 |
| Goat anti Mouse IgG (H+L) AlexaFluor 488 | ThermoFisher; A-11029 | IF | 1:1,000 |
| Goat anti Guinea Pig IgG (H+L) AlexaFluor 647 | ThermoFisher; A-21450 | IF | 1:1,000 |
| Goat anti Chicken IgY (H+L) AlexaFluor 647 | ThermoFisher; A-21449 | IF | 1:1,000 |

**Extended Data Table 4 | Software**

| Software | Source |
|---|---|
| FIJI/ImageJ v2.14 | https://imagej.net/software/fiji/ |
| Prism v10 | GraphPad |
| R v4.2.2 | https://www.r-project.org/ |
| Clampex v10.6 | Molecular Devices |
| Axon Multiclamp Commander v10.4 | Molecular Devices |
| Clampfit 10.7 | Molecular Devices |
| MATLAB R2021a | Mathworks |
| ImageLab Touch Software v1.0.0.15 | Bio-Rad |
| IGV browser v2.8.2 | UC San Diego and Broad Institute of MIT and Harvard |
| Minimap2 v2.26-r1175 | https://github.com/lh3/minimap2 |
| QIAxcelR | https://github.com/Delayed-Gitification/QIAxcelR |
| TapeStation Systems Software v3.2 | Agilent |
| QuantStudio Design & Analysis v1.5.2 | ThermoFisher Scientific |
| Maestro Pro – AxIS Navigator v3.7.2 | Axion Biosystems |
| Neural Metric Tool v4.0.5 | Axion Biosystems |
| Zen Blue v3.3 | Carl Zeiss AG |
